# BRIDGE: A Computational Workflow from Single Neurons to Network of Mean-Field Models

**DOI:** 10.64898/2026.07.31.742067

**Authors:** Ilaria Carannante, Damien Depannemaecker, Marmaduke Woodman, Pratik Purohit, Alain Destexhe

## Abstract

Mean-field models are extensively used in large-scale brain simulations because they provide a wieldy description of population dynamics while preserving key features of neural activity. Despite their widespread adoption, no common and reproducible methodology currently exists to systematically derive and validate mean-field models starting from biologically grounded single neuron dynamics. As a result, implementations are often ad hoc, difficult to reproduce and rarely reusable.

Here we introduce BRIDGE, a modular, open-source Python pipeline that enables the bottom-up reconstruction, analysis, validation, and simulation of mean-field models from single neurons. The framework integrates single neurons modelling, network simulations, extraction of population statistics, parameters analysis, quantitative comparisons between spiking neural networks and corresponding mean-field representations, and simulation of network of mean-fields. Its flexible architecture allows users to incorporate different neuron models and to generate region-specific or state-dependent mean-field formulations.

BRIDGE provides a reproducible foundation for developing biologically informed mean-field models suitable for large-scale and whole-brain simulations, supporting the transition from generic homogeneous population models toward region-specific ones.

**Graphical abstract:** 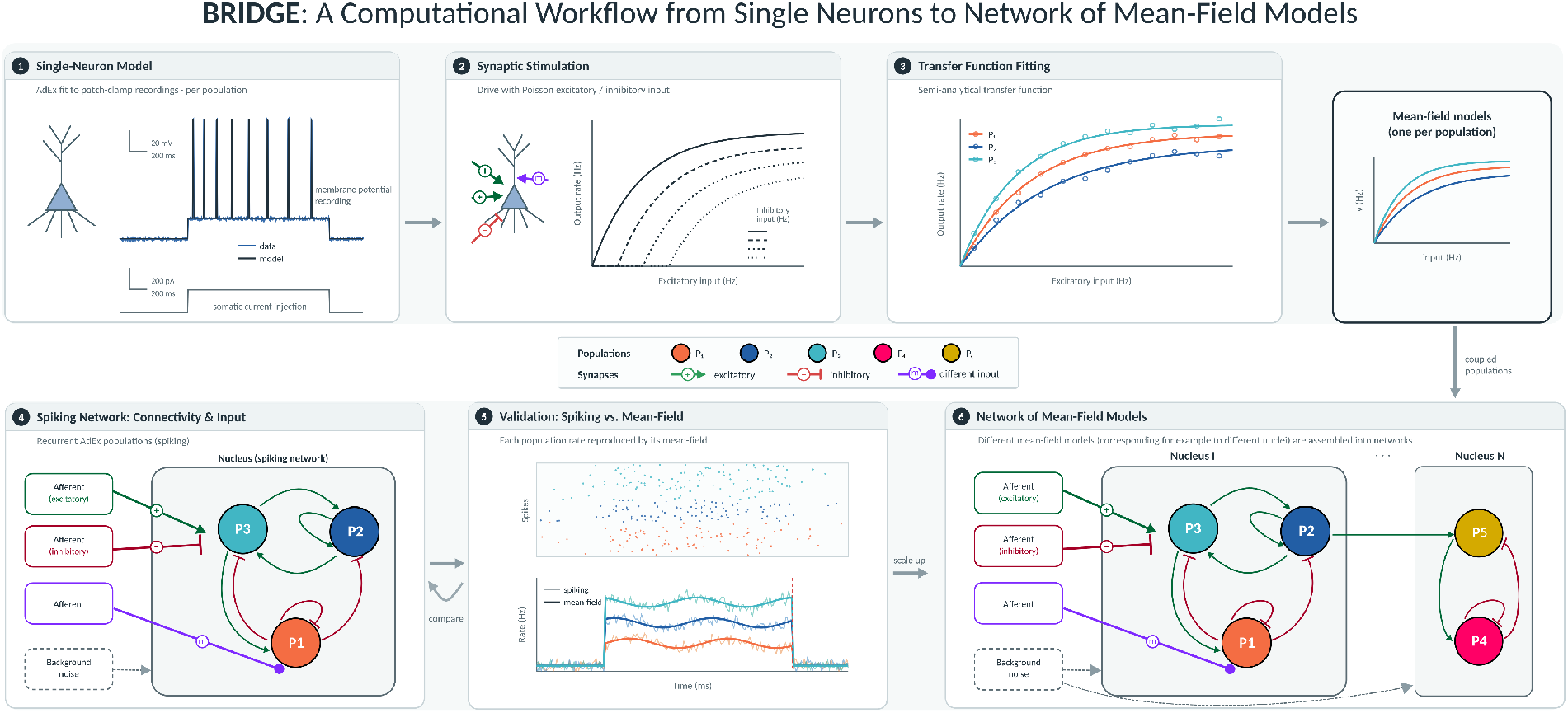

## 1 Introduction

Understanding how the coordinated activity of billions of neurons gives rise to brain function requires models that span multiple spatial scales. While detailed (multicompartmental or point-neuron) spiking neural network models capture the dynamics of individual neurons and microcircuits, simulating them at the scale of the whole brain remains computationally prohibitive.

Population-level descriptions, as mean-field or neural mass models, address this challenge by describing the collective dynamics of large neuronal populations through a small number of macroscopic variables, such as the mean firing rate (Wilson and Cowan [1972], Jansen and Rit [1995], Deco et al. [2008]). Over the past decades, these models have become a building block of large-scale and whole-brain simulation frameworks (Sanz Leon et al. [2013], Cakan et al. [2023]), where they are coupled through anatomical connectomes to reproduce macroscopic neuroimaging signals such as EEG, MEG, and fMRI (Deco et al. [2008], Sanz Leon et al. [2013]).

The central idea of the mean-field approach is to reduce a high-dimensional network of interacting spiking neurons to a low-dimensional system of ordinary differential equations describing collective population dynamics (El Boustani and Destexhe [2009], Brunel [2000]). By exploiting the statistical regularities of the asynchronous irregular regime that characterizes cortical activity *in vivo*, such reduction yields closed-form or semi-analytic expressions for the evolution of population firing rates and their fluctuations (El Boustani and Destexhe [2009], Zerlaut et al. [2018], Di Volo et al. [2019]). The resulting gain in scalability and analytical tractability comes at a cost of microscopic detail: single-neuron heterogeneity and precise spike timing.

Despite the maturity of the underlying theory and the widespread adoption of these models, deriving a mean-field model remains a largely manual, case-by-base process. Each new formulation requires substantial analytical and numerical effort typically involving fitting of the single-neuron models on electrophysiological data, computing the neuronal transfer function, and validating the reduced model against the full network (Zerlaut et al. [2018], Di Volo et al. [2019]). These steps are usually implemented ad hoc, with heterogeneous code bases, inconsistent conventions, and limited documentation. The consequences mirror a broader reproducibility concern in computational neuroscience, where published models are often difficult to reproduce independently and the associated code (when available) is rarely reusable beyond the study that produced it (Miłkowski et al. [2018]).

A further consideration relates the widespread use of generic population models in whole-brain simulations. For simplicity, many frameworks assign the same mean-field model to several, if not all, brain regions, effectively treating the brain as a network of identical, homogeneous units. However, brain regions differ in their cellular composition, microcircuit organization, and connectivity, and these local properties shape regional dynamics in a way that generic population models can not capture (Lorenzi et al. [2025]). There is therefore growing interest in region-specific mean-field models derived from biologically grounded single-neuron and microcircuit descriptions.

Here we address this methodological gap by introducing BRIDGE (Bottom-up Reconstruction and Inference of Dynamics in Generalized Ensembles), a modular, open-source Python pipeline for the bottom-up reconstruction, analysis, validation, and simulation of mean-field models from single neurons. The pipeline integrates the full workflow within a single reproducible framework: single-neuron modeling, spiking network simulation, extraction of population statistics, parameter analysis, quantitative comparison between the spiking network and its mean-field representation, and simulations of network of mean-fields. Its modular architecture lets users substitute different neuron models and generate region-specific or state-dependent formulations (for example pathological states) without re-implementing the derivation from scratch. The pipeline provides a reproducible foundation for building biologically informed population models, supporting the transition from generic homogeneous models towards region-specific formulations suitable for large-scale and whole-brain simulations.

## 2 Methods

In this section we describe the *bottom-up steps* of the workflow.

All simulations were performed in Python with the support of the simulator `Brian 2` (Stimberg et al. [2019]). Functions provided by `Brian 2` are referred to using the notation `b2.func_name`. Implementation details and analysis are documented in the accompanying GitHub repository.

### 2.1 Single Neuron Model

We use the Adaptive Exponential Integrate-and-Fire model (AdEx), introduced by Brette and Gerstner [2005], and analysed by Touboul and Brette [2008]. The AdEx model provides a good compromise between accurately capturing the rich and diverse neuronal dynamics and maintaining computational efficiency, making it particularly suitable for large-scale network simulations. Importantly, AdEx is also well suited for studying neuromodulatory effects (Guarino et al. [2025]), and its parameters retain a clear biological interpretation.

In this framework, each neuron is represented as a single compartment, and its membrane potential *V* (*t*) and adaptation current *w*(*t*) evolve according to two coupled differential equations:

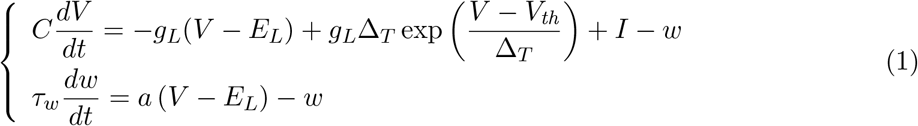

with *C* membrane capacitance, *g*_*L*_ leak conductance, *E*_*L*_ leak equilibrium potential, Δ_*T*_ exponential slope factor, *V*_*th*_ threshold potential, *I* input currents, *τ*_*w*_ adaptation time constant, and *a* subthreshold adaptation conductance.

When the membrane potential increases, the exponential term generates an upswing in the trajectory, leading to the initiation of an action potential. At spike detection (*V*_*peak_detect*_), reset conditions are triggered:

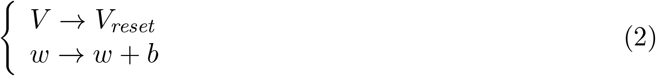

hence the voltage is reset to *V*_*reset*_, and *w* is increased by *b*, the spike-triggered adaptation.

In this study, we consider three neuronal populations: inhibitory fast spiking neurons (FS) and excitatory regular spiking neurons (RS), the latter in both an adaptive (*b* ≠ 0) and a non-adaptive (*b* = 0) configuration. The parameter values used are reported in Table 1. From these, we construct two networks: one combining FS and non-adaptive RS neurons, and one combining FS and adaptive RS neurons.

**Table 1:** Parameters of the FS and RS neuron models adapted from Depannemaecker et al. [2025].

| Parameters | FS | RS | Unit |
| --- | --- | --- | --- |
| $C_m$ | 0.2 | 0.2 | nF |
| $g_L$ | 0.01 | 0.01 | $\mu S$ |
| $E_L$ | -65 | -65 | mV |
| $I_e$ | 0 | 0 | nA |
| $a$ | 0 | 0 | nS |
| $b$ | 0 | 0.1 | nA |
| $\tau_w$ | 1 | 500 | ms |
| $V_{th}$ | -48 | -50 | mV |
| $\Delta_T$ | 0.5 | 2 | mV |
| $V_{reset}$ | -65 | -65 | mV |
| $V_{peak\_detect}$ | -47.5 | -40 | mV |
| $t_{ref}$ | 5 | 5 | ms |

To derive AdEx models of neuronal populations, the Python pipeline developed by Guarino et al. [2025] can be applied. Indeed, starting from elecrophysiological data and following the different steps, the user can extract the characteristic features of the voltage recordings and initiate a parameter investigation that leads to (new) population specific AdEx models. In addition, the authors also provide a large dataset of developed models for multiple brain region both in control conditions and under different neuromodulators.

We incorporate and expand such framework by guiding the users to the construction of neural network, derivation of mean-field models, their analysis, and simulations.

#### Implementation

Model parameters, initial conditions and their units are specified in structured . `json` configuration files. The model parameters can be supplied directly when they are already known, or obtained by running the parameter-search stage described above. The AdEx equations (Eq. 1) are explicitly defined and passed to `b2.NeuronGroup`, together with the corresponding threshold and reset conditions (Eq. 2). At runtime, the simulator integrated the dynamical systems, and the voltage *V* (*t*) and adaptive current *w*(*t*) are recorded via `b2.StateMonitor` for subsequent analysis and visualization.

Alternative single neuron models can be implemented by providing the corresponding equations and reset conditions. As a worked example of this modularity, we include the extended generalized leaky integrate-and-fire (E-GLIF) model (Geminiani et al. [2018], see Supplementary Material).

### 2.2 Synaptic Model

Neuronal communication is mediated by synaptic transmission. Although it involves several receptor types, computational models of neural networks commonly reduce this diversity to a single excitatory and a single inhibitory channel, which we model as conductance-based. In this case, the input current *I* (in Eq. 1) is defined as the sum of an excitatory and an inhibitory synaptic current:

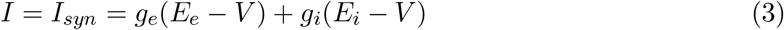

where *g*_*e*_ and *g*_*i*_ are the excitatory and inhibitory conductances, respectively, and *E*_*e*_ and *E*_*i*_ are the corresponding reversal potentials. The conductances follow first-order kinetics, decaying exponentially between presynaptic events and increasing by a fixed quantal amplitude upon each incoming spike:

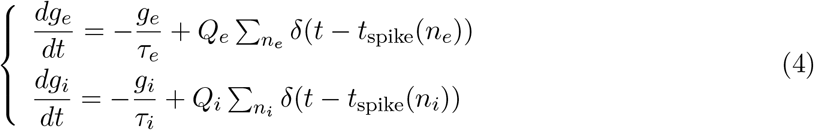

where *Q*_*e*_ and *Q*_*i*_ are the excitatory and inhibitory quantal conductances, respectively; *n*_*e*_ and *n*_*i*_ index the number of incoming excitatory and inhibitory presynaptic synapses; *t*_*spike*_(*n*_*e*_) and *t*_*spike*_(*n*_*i*_) denote their spike arrival times; and finally *τ*_*e*_ and *τ*_*i*_ are the synaptic decay time constants. The values used in this study are reported in Table 2.

**Table 2:** Excitatory and inhibitory synaptic parameters (from Depannemaecker et al. [2025]).

| Synaptic parameters |  | Unit |
| --- | --- | --- |
| $E_e$ | 0 | mV |
| $Q_e$ | 1.5 | nS |
| $n_E$ | 8000 | — |
| $E_i$ | -80 | mV |
| $Q_i$ | 5.0 | nS |
| $n_I$ | 2000 | — |
| $p$ | 0.05 | — |
| $\tau_{\text{syn}}$ | 5 | ms |

This description generalizes to an arbitrary set of synaptic channels {*s* }, each characterized by a reversal potential *E*_*s*_, a quantal conductance *Q*_*s*_, and a decay time constant *τ*_*s*_. In this more detailed case, distinct synapses are described with their own dynamics rather than grouped into a single excitatory and a single inhibitory channel. The synaptic current becomes

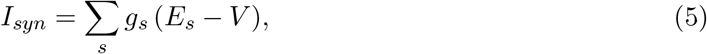

and the conductances evolve as

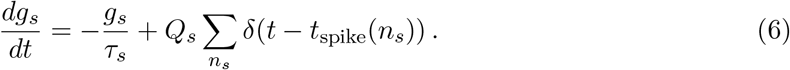

The implemented framework accommodates an arbitrary number of synaptic channels, as well as alternative formulations depending on the modeled brain region. In the E-GLIF example, the synapses are indeed modeled with alpha-function conductances and the modeled neuron receives three distinct synaptic channels (see Supplementary Material).

#### Implementation

Synaptic parameters, connection probabilities, and input rates are specified in .json configuration files. The conductance dynamics (Eq. 6) are explicitly defined within the neuronal equations and integrated together with the membrane dynamics (through Eq. 5). Synaptic inputs arise from two sources: intranuclear (i.e., synapses between the defined neuronal populations) and external. Intranuclear synapses are instantiated using `b2.Synapses` between neuronal populations (see Neural Network section) and the presynaptic spike events induce discrete updates of the corresponding conductances via the specified quantal amplitudes. External drive is first generated using `b2.PoissonGroup`, with input population sizes and firing rates specified in the configuration files, and the synapses from this “{external population”{ are then instantiated as in the intranuclear case.

### 2.3 Neural Network

We constructed two neural networks containing 10 000 neurons, divided into two populations, fast-spiking inhibitory neurons (FS, 20%) and regular-spiking excitatory neurons (RS, 80%). In one network configuration, RS neurons include spike-triggered adaptation; in the second configuration, adaptation is removed (*b* = 0). The two configurations share identical connectivity and external drive, allowing direct comparison between adaptive and non-adaptive dynamics.

Neurons are randomly connected both within and across populations with a fixed connection probability of 5%. Since the networks do not exhibit self-sustained activity under the chosen parameter regime, they require external drive.

#### Implementation

Network composition, connection probabilities, and external input specifications are collected in `.json` configuration files. These include the total number of neurons, the relative size of each population, recurrent connection probabilities, the type and number of external inputs, and their firing rates.

Neuronal populations are instantiated using the same single-neuron construction described above. Intranuclear synapses are then created between and within populations using `b2.Synapses`, with connections drawn according to the specified probability. Synaptic type is determined by the presynaptic population, with RS and FS forming excitatory and inhibitory synapses, respectively. Upon a presynaptic spike, the synapse increments the corresponding conductance by the discrete amplitude *Q*_*s*_ (as described in Eq. 6).

External drive is generated using `b2.PoissonGroup`, which defines presynaptic populations emitting stochastic spike trains with specified rated. Synaptic projections from these populations to the network are instantiated using `b2.Synapses`, following the same conductance-based update rule as before.

State and spike monitors are defined to record membrane potentials and spike times during simulation. From these recordings, raster plots and population firing rates (mean and standard deviation, computed with given bin size) are obtained. Simulations outputs can optionally be stored in `.h5` files for further analysis.

### 2.4 Mean-field

The mean-field (MF) model provides a low-dimensional description of the network, replacing the thousands of coupled single-neuron equations with a small system of ordinary differential equations governing the mean firing rate of each population. Following the formalism of Zerlaut et al. [2018] and Di Volo et al. [2019], the core element of this reduction is the *transfer function* of each population, i.e. the stationary output firing rate of a neuron as a function of its (excitatory and inhibitory) input rates. We first describe how the transfer function is characterized, fitted, and analyzed, and then how the population-specific transfer functions are assembled into the population rate equations.

#### 2.4.1 Transfer Function

The transfer function maps the mean presynaptic input rates onto the stationary output firing rate of the postsynaptic population. In the most common case, as anticipated, the drive is grouped into a single excitatory (*ν*_*e*_), and a single inhibitory (*ν*_*i*_) channel:

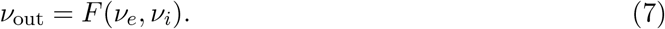

However, more generally, the input can come from several distinct channels, for instance multiple excitatory or inhibitory channels with different synaptic time constants or quantal conductances, in which case the transfer function depends on the full set of input rates (*ν*_1_, …, *ν*_*n*_).

We adopt the semi-analytical form of Zerlaut et al. [2018], in which the output rate is expressed through the statistics of the subthreshold membrane potential and a phenomenological effective threshold. Under the assumption of asynchronous irregular activity, each channel is treated as shot noise, giving the mean *µ*_*V*_, standard deviation *σ*_*V*_, and autocorrelation time *τ*_*V*_ of the membrane potential:

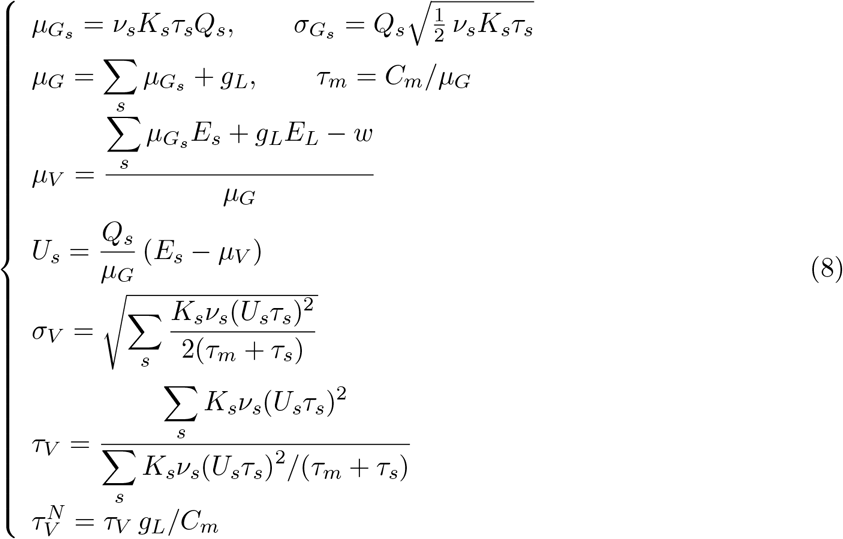

where the index *s* runs over all input channels, each characterized by its mean rate *ν*_*s*_, number of afferent connections *K*_*s*_, quantal conductance *Q*_*s*_, synaptic decay time constant *τ*_*s*_, and reversal potential *E*_*s*_; *g*_*L*_, *E*_*L*_, and *C*_*m*_ are the leak conductance, leak reversal potential, and membrane capacitance, and *w* is the adaptation current (measured directly during data generation as the mean adaptation variable over the output-rate window, and set to zero in the non-adaptive case). Each channel contributes to the total conductance *µ*_*G*_ and, through its reversal potential *E*_*s*_, to the mean membrane potential *µ*_*V*_ .

The output firing rate is then:

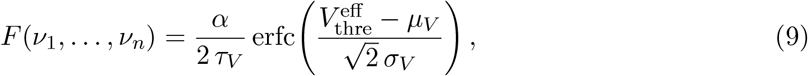

with erfc the complementary error function and *α* a global scaling factor (Carlu et al. [2020]). The effective threshold is expressed as a second-order polynomial in the normalized membrane potential statistics:

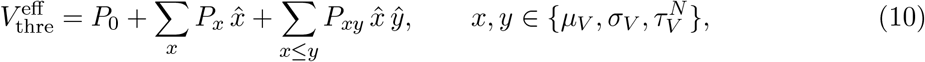

where 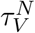 is the autocorrelation time normalized by the leak (resting) membrane time constant (*C*_*m*_*/g*_*L*_), each 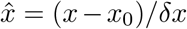, and {*P*_0_, *P*_*x*_, *P*_*xy*_} is a set of ten coefficients determined by fitting.

#### Data generation

The transfer function is characterized empirically by simulating a population of *N* uncoupled neurons of a single type (default value is *N* = 50), driven by independent Poisson inputs. The neurons are not recurrently connected, so that each one independently samples the input-output relationship, and averaging across the population gives the mean output rate *ν*_out_ and its standard deviation *σ*_out_ at each input combination. The input space is sampled on a grid over the channel rates: once the range and step of each *ν*_*s*_ are fixed, every combination is simulated to extract *ν*_out_ and *σ*_out_. In the case of one excitatory and one inhibitory input (*s*∈{ *e, i* }), this results in a matrix (*ν*_*e*_, *ν*_*i*_, *ν*_out_, *σ*_out_). The analytical transfer function (Eq. 9) is then fitted to these data.

#### Fitting

The ten polynomial coefficients and the scaling factor *α* are estimated from the simulated data through a two-stage procedure nested within a search over *α*.

For a given *α*: (i) Eq. 9 is inverted to obtain, at each grid point, an empirical effective threshold

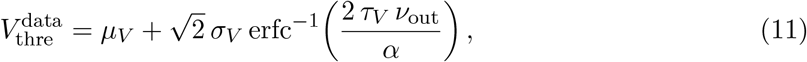

and the coefficients are fitted in threshold space by minimizing the squared error (SLSQP), providing an initialization; (ii) the coefficients are refined by directly minimizing the mean squared error (Nelder–Mead) between the measured output rates and the transfer function prediction. Both minimizations use `scipy.optimize.minimize`. This two-stage fit is repeated over a user-defined range [*α*_min_, *α*_max_] with fixed *α*_step_ (default 0.1–2.0, step 0.001), and the pairs (*α, P* ) with the lowest rate-space error are retained and saved. The procedure is applied independently to each population.

### 2.5 Parameter investigation

The fitting procedure returns the *N* = 10 parameter sets of lowest mean error. Reporting only the best of these discards what the ensemble records: whether the retained sets sample one solution repeatedly or several distinct solutions of comparable quality.

Individual coefficients are compared through their coefficient of variation, CV_*i*_ = | *σ*_*i*_*/µ*_*i*_|, taken across the retained sets. Their joint structure is characterized by the Pearson correlation matrix and by a principal component analysis of the standardized coefficients. Specifically, the standardization is applied so that the decomposition describes co-variation rather than the relative scale of coefficients.

Discrete structure in the ensemble is detected by Ward clustering of the standardized coefficients, the number of clusters being selected by silhouette score over *k* ∈ [2, 4] and set to one when no partition reaches a silhouette of 0.55. When more than one cluster is found, the retained sets are not independent draws

The consequence of the coefficient spread for the fitted surface is assessed directly. All retained sets are evaluated on the input grid (*ν*_1_, …, *ν*_*n*_) and, at each grid point, the standard deviation of the predictions across the ensemble is compared with the standard deviation of the simulated firing rate. Where the former falls below the latter, the retained sets are not distinguishable by the data and their coefficient spread does not constitute an uncertainty on the transfer function. The absolute and relative deviation of the best fit from the simulated rate are mapped over the same grid to delimit the domain over which the fitted transfer function is accurate.

Populations are compared in a common basis. The *M* ensembles are stacked into a single matrix *X* ∈ ℝ ^(*MN* )×*np*^, whose columns are standardized using the mean and standard deviation of the pooled matrix, *Z* = (*X* − *µ*_pool_)*/σ*_pool_. A single principal component analysis is fitted to *Z*, generating a component matrix *W*, and each population is projected onto it as *S*_*m*_ = *Z*_*m*_*W* ^T^, where *Z*_*m*_ denotes the rows of *Z* belonging to population *m*. Since the sign of a principal component is not determined by the decomposition, the components are oriented against a reference population before their loadings are compared. Each component of the reference population is oriented so that its largest magnitude loading is positive, and the corresponding component of every other population is multiplied by sign 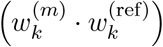 . The convention is well defined only where the two components are not close to orthogonal, so the inner products are reported alongside the comparison and components for which 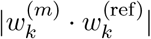 is small are notcompared. The analysis is implemented in `scikit-learn` and available as a notebook that is applied unchanged to each population.

### 2.6 Mean-field validation against spiking networks

Each reconstructed MF model was validated against the spiking network (NN). Given a reconstructed model, BRIDGE quantifies, across the input space, where the MF reproduces the NN and it localizes the origin of any discrepancy. The user can then establish an explicit regime of validity of their own model.

Each population received independent external excitatory and inhibitory drive from Poisson populations (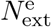 and 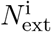 units, connected to the target population with probability *p*_ext_). The instantaneous rate shared by the units of each external Poisson population was set by a rectified Ornstein-Uhlenbeck process (*τ*_OU_ = 50 ms, *σ*_OU_ = 2) whose mean was fixed to the external drive *ν*_exc_ or *ν*_inh_; the two means were varied independently on a regular grid over [0, 30] Hz. The MF was integrated under the matched external drive (the same mean afferent rates entering its transfer function) and analyzed over the same window, so that the two descriptions are compared under identical simulations.

BRIDGE characterizes the agreement in two complementary ways (Figure 6). The *closed-loop* comparison evaluates the full MF as a self-consistent dynamical system, in which each population’s recurrent input depends on its own output rate, and compares its stationary rate to the network (*F*_MF_*/F*_NN_). The *open-loop* comparison instead isolated the transfer function: at each grid point we take the network’s measured rates, reconstruct the afferent input each population received (external drive plus the recurrent contribution implied by the measured rates, following the network connectivity), and evaluate the transfer function once at that operating point, comparing its prediction to the measured rate. Together these separate the two possible sources of mismatch: an inaccurate transfer function versus amplification of a transfer-function residual by the recurrent loop. This is the practical diagnostic the pipeline exposes: it tells a user whether an observed mismatch should be addressed by refitting the transfer function or is an intrinsic, regime-dependent property of the recurrent network, and therefore how to act on it. BRIDGE summarises accuracy at each operating point by the relative error |*F*_MF_*/F*_NN_ − 1| and classifies points as in quantitative ( ≤ 15%) or qualitative ( ≤ 50%) agreement; operating points at which the network rate falls below 0.5 Hz are excluded, as the ration is then ill-defined. To relate the regime of validity to the assumptions of the semi-analytical formalism, BRIDE also computes the membrane-potential fluctuation amplitude *σ*_*V*_ at each operating point, which distinguishes the fluctuation-driven regime in which the transfer function is derived from the mean-drive regime in which it degrades.

#### Implementation

For each external input rate grid point, the network (see Neural Network section) is driven by external `b2.PoissonGroup` populations whose shared instantaneous rate is a rectified Ornstein-Uhlenbeck process with mean set to the grid value. The rate is supplied as a `b2.TimedArray` and set to zero outside the stimulation interval. Projections from these external populations follow the same conductance-based update as the intranuclear synapses (Eq. 6). Spikes are recorded with `b2.SpikeMonitor`, and population firing rates (mean and standard deviation, at a given bin size) are computed over the steady-state window, and stored in .h5 files. The corresponding MF model is integrated with `scipy.integrate.solve_ivp` under the matched external drive and analysed over the same window, and its stationary rates are stored in the same format.

The two descriptions are then compared point by point. The closed-loop agreement is the ration of the mean-field to the network stationary rate. The open-loop agreement is obtained by evaluating the transfer function Eq. 9) at the operating point measured in the network (the external drive together with the recurrent input implied by the measured rates) and comparing its prediction to the network rate. The membrane-potential fluctuation amplitude *σ*_*V*_ is evaluated at the same operating points.

### 2.7 Phase-plane analysis of the mean-field model

Once the transfer functions have been fitted for each population, the assembled network admits a low-dimensional description whose dynamics can be studied directly in the plane. BRIDGE analyze the populations through their mean firing rates (*ν*_FS_ and *ν*_RS_), and characterize the resulting two-dimensional system by its nullclines, fixed points, and their dependence on external drive and model parameters.

#### Reduce dynamical system

In the first-order approximation, the two population rates *ν* = (*ν*_FS_, *ν*_RS_) are governed by:

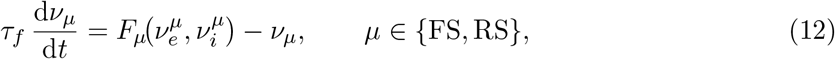

where *F*_*µ*_ is the semi-analytical transfer function of population *µ* and *τ*_*f*_ the population time constant. The presynaptic identity sets the synapse type: RS is excitatory and FS is inhibitory, so the excitatory and inhibitory drives seen by each population combine the recurrent rate of the corresponding presynaptic population with the external input. For population *µ*, the effective input rates entering *F*_*µ*_ are:

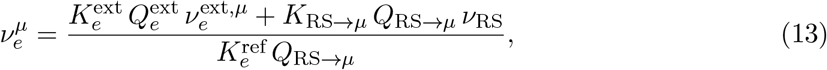

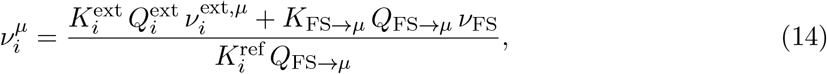

where *K*_*η → µ*_ = *p N*_*η*_ are the mean recurrent in-degrees implied by the connection probability *p* and the presynaptic population size 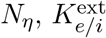 the external in-degrees, and *Q* the corresponding quantal conductances, consistence with the channel quantities *K*_*s*_, *Q*_*s*_ of Eq. 8. The normalization by the reference in-degree and conductance 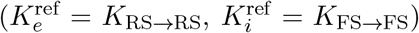 expresses the total synaptic drive as an equivalent presynaptic rate through the reference pathway, matching the input convention used during transfer-function fitting. The external rates 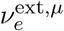 and 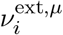 are the four control inputs of the reduces system; their default values are read from the `rates` block of the network configuration, so the analysis starts at the operating point at which the network was specified.

#### Nullclines and fixed points

The *ν*_*µ*_-nullcline is defined by d*ν*_*µ*_*/*d*t* = 0, i.e. *F*_*µ*_(*ν*) = *ν*_*µ*_. The two nullclines are traces as the zero level sets of the right-hand side of Eq. (12) on a regular grid over the displayed rate window. Their intersection are the fixed points *ν*^∗^ of the network. Fixed points are located by damped Newton iteration (`scipy.optimize.fsolve`) seeded from teh grid nodes, retaining converged solutions within the analysis window whose residual ∥*F* (*ν*^∗^) −*ν*^∗^∥ falls below a fixed tolerance and dedupicating solutions that coincide to within a rate tolerance.

#### Linear stability

Each fixed point is classified from the eigenvalues of the Jacobian of Eq. (12):

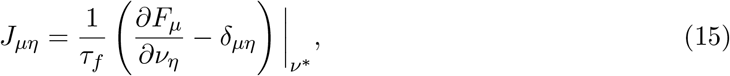

evaluated by finite differences, where the dependence of *F*_*µ*_ on *ν*_*η*_ runs through the effective inputs 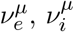 (Eqs. 13–14). A fixed point is a stable node or focus when both eigenvalues have negative real part (focus if the imaginary parts are non-zero), and unstable node or focus when both have positive real part, and a saddle when the real parts have opposite sign.

#### Trajectories and flow

Trajectories are obtained by integrating Eq. (12) from a chosen initial condition with an explicit adaptive Runge-Kutta scheme (`scipy.integrate.solve_ivp`, RK45), and are displayed both in the (*ν*_FS_, *ν*_RS_) plane and as rate time courses. The direction field is sampled on a grid to visualize the flow toward the stable state.

#### Parameter continuation

The organization of fixed points are a function of a single control parameter (an external input rate, the population time constant, or the transfer function scaling factor *α*) is obtained by sweeping that parameter over a range and recomputing the fixed points and their stability at each step. To follow branches robustly across the sweep, the fixed points found at one parameter values seed the Newton search at the next, so continuous branches are tracked by continuation. The resulting bifurcation diagram plots the fixed-point rates against the control parameter, colour-coded by stability, and exposes the appearance and disappearance of solutions (saddle-node transitions) as the drive is varied.

#### Implementation

The phase-plane analysis is implemented as an interactive component of BRIDGE. Nullclines, fixed points, trajectories, and parameter sweeps update in response to the external drive and model parameters, and each panel can be exported as a vector graphics.

### 2.8 Network of Mean-Fields

Brain function is widely understood to emerge from interactions among neuronal assemblies, so beyond the single-node model we study the network-level dynamics of coupled nodes. Each node represents a neuronal assembly and groups one or more neuronal populations. The connectivity between nodes can be defined from several sources (user-defined weights based on experimental data, structural connectivity from tractography or tracing, functional connectivity), and enters the model through three per-connection quantities: the synaptic receptor type, a connection weight, and a propagation delay.

The network is described at population level. Populations are indexed by *µ* and inheriting the corresponding transfer function *F*_*µ*_ together with the single-cell parameters. The mean firing rate *ν*_*µ*_ of population *µ* is described by

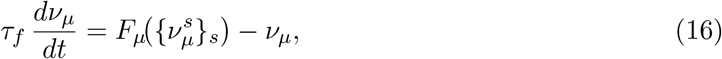

where *τ*_*f*_ is the population rate time constant and *F*_*µ*_ is evaluated on the effective afferent rate 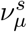 on each synaptic channel *s*.

The channel *s* afferent rate collects the delayed activity of every population presynaptic to *µ* on that channel, together with an (optional) external stimulus:

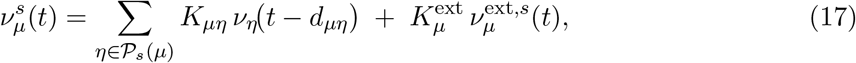

where *P*_*s*_(*µ*) is the set of populations presynaptic to *µ* on channel *s, K*_*µη*_ = *N*_*η*_ *p*_*µη*_ *w*_*µη*_ is the mean in-degree (presynaptic size *N*_*η*_, connection probability *p*_*µη*_, connection weight *w*_*µη*_), and *d*_*µη*_ is the propagation delay specified for that connection (which can be seen as the ratio between the physical distance between nodes and the conduction velocity). Connections can be intra- or inter-node, so recurrent and long-range projections are handled uniformly. The external stimulus enters through its own in-degree 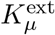 and imposed rate 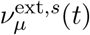, providing a means of *in silico* stimulation of any chosen population.

#### Implementation

The network layer is built directly on the transfer-function procedure (Section 2.4.1): the transfer function is evaluated for every population at every step. A network is specified by a configuration file containing: (i) the population name, cell type, node, initial rate, and (ii) the per-type single-cell parameters together with the transfer function coefficients. No intermediate fitting output or simulation data is read at run time, so a network is fully reproducible from its configuration alone.

The representation is deliberately general. A node may contain an arbitrary number of populations of arbitrary types, and connections may be intra- or inter-node, so networks whose nodes represent different regions with distinct population composition can be easily assembled (provided the required parameters). Each population may additionally receive user-defined external stimuli. The simulation output is the time course of every population’s rate, from which node activity, responses to stimuli, and inter-node propagation are analyzed and visualized.

## 3 Results

BRIDGE, the pipeline presented in this work, reconstructs mean-field models through a sequence of modular steps: single-neuron and synaptic modeling, neural network dynamics, fitting of the transfer function, analysis of the fitted parameters, mean-field validation and network of mean-fields simulations.

Throughout this section we demonstrate the workflow using AdEx neurons with conductance-based synapses, however in the Supplementary Material we show that BRIDGE applies equally to E-GLIF neuron model with alpha-function synapses.

### 3.1 Single neuron models

The first stage of BRIDGE consists of defining the single-neuron models that will be used to build the spiking network and mean-field. It supports two complementary workflows. When no suitable single-neuron model is available for the neuronal population of interest, users can first derive population-specific AdEx parameters directly from electrophysiological recordings using the parameter-search framework of Guarino et al. [2025] (integrated within BRIDGE, see Methods). On the other hand, when validated parameters are already available from previous studies or model repositories, they can be imported and used for simulation and visualization. Figure 1 illustrates this process for the three neuronal populations used throughout this work (parameters in Table 1). Fast-spiking (FS) neuron exhibits sustained high-frequency firing during the stimulus. In contrast the regular-spiking (RS) neuron displays pronounced spike-frequency adaptation, as the adaptation current accumulates, successive interspike intervals become progressively longer, reducing the rate over time, while the prolonged after-hyperpolarization following offset reflects the slow decay of the adaptation variable. Removing the spike-triggered adaptation (RS_no_adapt) restores sustained firing, producing a response that closely resembles that of the FS neuron despite their different intrinsic membrane properties.

**Figure 1.**
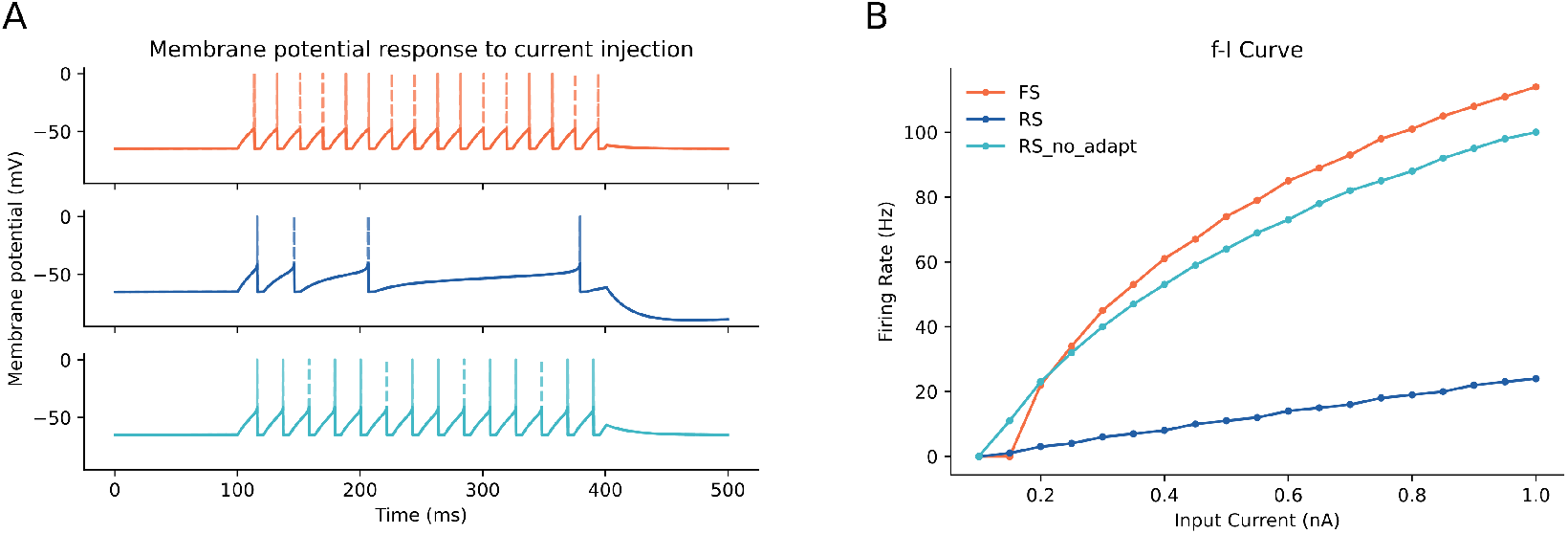
Single-neuron dynamics. A) Membrane potentials response to a step current injection *I* = 0.3 nA applied from *t* = 100 ms to *t* = 400 ms. Top (orange): fast-spiking (FS) neuron, characterized by a short membrane time constant and the absence of adaptation (*a* = *b* = 0), which sustains hight-frequency, regular discharge for the entire duration of the stimulus. Middle (blue): regular-spiking (RS) neuron, whose adaptation current (*b* = 0.1 nA and *τ*_*w*_ = 500 ms) produces a marked spike-frequency adaptation: the interspike interval lengthens progressively and the after-hyperpolarisation persisting beyond stimulus offset reflects the slow decay of *w*. Bottom (teal): identical to the regular-spiking neuron except that the adaptation is removed (RS_no_adapt). B) Stationary firing rate as a function og the injected current (*f* -*I* curve) for the three neurons. The FS and RS_no_adapt curves are close to each other over the whole range, with the FS reaching higher rates at large currents.

The stationary *f* -*I* curves (Figure 1B) place FS and RS_no_adapt close together across the current range, with FS reaching higher rates at large currents, while adaptation lowers and saturates the RS response.

### 3.2 Conductance-based synaptic model

We next inspect the conductance-based synaptic model (independently of the recurrent network, Figure 2A, parameters in Table 2). A single presynaptic spike produces excitatory and inhibitory currents, whose waveforms are set by the decay constants (in this case it is the same value for both synaptic models). The peak of the postsynaptic current is linear in the holding potential, with slope set by the quantal conductance and zero crossing at the channel reversal potential (Figure 2B), so the polarity of a synaptic event follows the reversal potential relative to the membrane. When driven by stochastic Poisson inputs, the three neurons respond with membrane potential fluctuations around their resting state, while transient increases in excitatory or inhibitory input selectively depolarized or hyperpolarized the membrane (Figure 2C).

**Figure 2.**
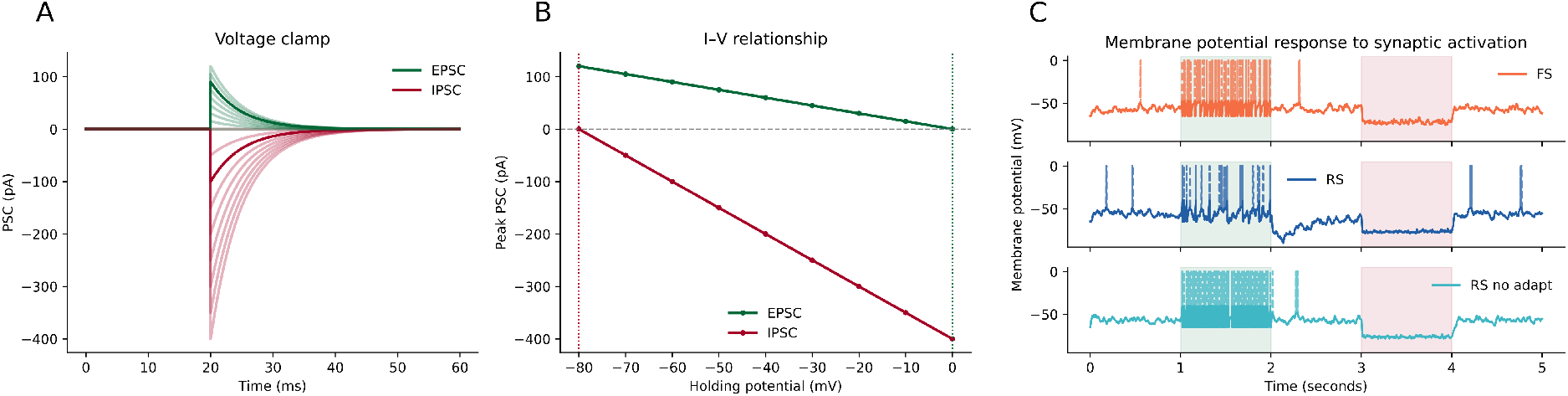
Fig 2. Conductance-based synaptic model. A) Voltage-clamp response to a single presynaptic spike derive at *t* = 20 ms, for holding potentials from − 80 to 0 mV in 10 mV steps (light traces); the dark traces correspond to a holding voltage of − 60 mV. Excitatory (green) and inhibitory (red) currents share the same waveform, set by the decay constants (which in this case are equal *τ*_*e*_ = *τ*_*i*_ = 5 ms), and differ in amplitude and in the sign of the driving force. B) Peak postsynaptic current as a function of holding potential. The relation is linear, its slope given by the quantal conductance (*Q*_*e*_ = 1.5 nS and *Q*_*i*_ = 5 nS) and its intercept the the corresponding reversal potential. The polarity of a synaptic event is determined by the reversal potential of the channel relative to the membrane potential. C) Membrane potential of each neuron during a 5 s simulation in which the cell receives Poisson background input (0.5 Hz), an additional excitatory drive between *t* = 1 and *t* = 2 s (green shading) and inhibitory drive between *t* = 3 and *t* = 4 s (red shading).

### 3.3 Network dynamics emerge from recurrent interactions

BRIDGE next step is to assemble the single-neuron and synaptic components into recurrent networks. This specif ones contain the inhibitory FS population and the excitatory RS populations. They are connected both within and across populations, so that an external input targeting one population could influence the other through the recurrent circuitry. To isolate the contribution of spike-frequency adaptation, we compared two otherwise identical networks: one containing adaptive RS neurons and one in which spike-triggered adaptation was removed.

In the adaptive FS-RS network (Figure 3 A-C), excitation delivered directly to the FS population increased FS activity without recruiting the RS population. Since FS neurons provide inhibitory output, their activation maintained the RS population near silence. In contrast, excitation of the RS population recruited both populations. The externally driven increase in the RS firing activated FS neurons through recurrent excitatory projections producing a coordinated response of the two populations. Thus, although the external stimulation targeted only the excitatory population, recurrent interactions distributed the response across the network.

**Figure 3.**
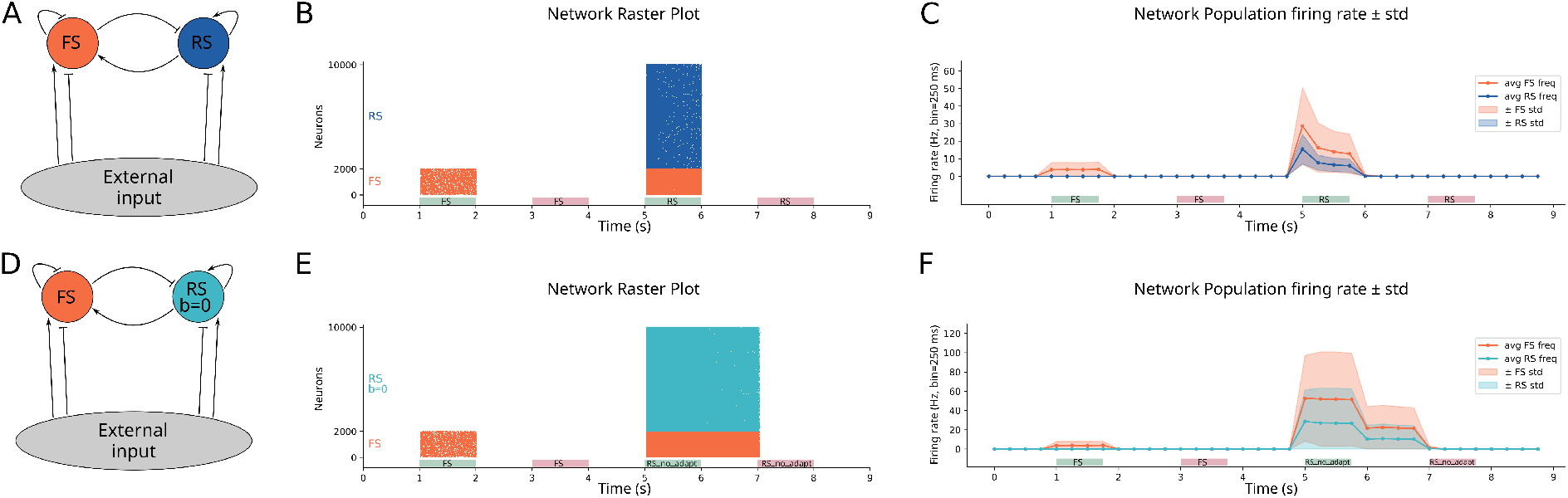
Spiking network response to targeted external drive, with and without adaptation. Both networks contain *N* =10 000 neurons (20 % FS, 80 % RS) randomly connected with probability *p* = 0.05. Within a stimulation window, each afferent fires at 2 Hz (excitatory drive) or 1 Hz (inhibitory drive), independently of the population targeted. The colored boxes below the raster and firing-rate plots indicate the stimulation windows and the population receiving the external input: green denotes excitatory stimulation, red represents inhibitory stimulation, and the label inside each box define the targeted popularion. A) Architecture of the first network, recurrently and mutually connected FS (inhibitory) and RS (excitatory, adaptive) populations, both receiving external excitatory and inhibitory drive. B) Raster plot of the FS–RS network. Excitation of the FS population (1–2 s) activates FS neurons and, the population being inhibitory, does not recruit RS neurons. Excitation of the RS population (5–6 s) engages both populations through the recurrent excitatory connections. The two inhibitory stimulation windows (3–4 s and 7–8 s) are applied while the network is already silent and produce no visible change. C) Population firing rate of the FS (orange) and RS (blue) populations for the simulation in B), computed in 250 ms bins. Shaded areas denote ± one standard deviation across the neurons of the population. Adaptation therefore both limits the evoked rate and prevents the recurrent excitation from sustaining itself. D) Architecture of the second network, identical to A) except that the excitatory population is non-adaptive (*b* = 0; RS_no_adapt). E) Raster plots of the FS–RS_no_adapt network under the same stimulation protocols as in B). F) Population firing rates for the simulations of E). Note the difference in ordinate scale between C) and F).

Removing the adaptation from the excitatory population substantially changed the network response (Figure 3 D-F). Under the same external stimulation protocol, the RS_no_adapt generated a much larger response, which in turn recruited stronger FS activity. The difference in firing-rate scale between the two networks shows that this effect does not arise from connectivity or external input, but from intrinsic dynamics of the excitatory neurons.

Spike-frequency adaptation therefore acts as a population-level negative feedback: it limits excitatory amplification, reduces the activity transmitted through the recurrent loop, and stabilizes the network response.

### 3.4 Transfer function characterization and fitting

The next step of BRIDGE is the reconstruction of the transfer function, which provides the link between microscopic spiking activity and mesoscopic population level descriptions. BRIDGE computes the stationary relationship between the mean presynaptic input rates and the mean output firing rate of a population.

In the present networks the input is grouped into a single excitatory and a single inhibitory channel, so this relationship is *ν*_out_ = *F* (*ν*_*e*_, *ν*_*i*_) (Eq. 7). However the formalism extends to an arbitrary set of synaptic channels, as described in the Methods. For instance, the E-GLIF (Golgi neuron) example in the Supplementary Material contains three distinct channels (two distinct excitatory and one inhibitory, Supplementary Figure 2).

The transfer function is the closure that allows the high-dimensional NN to be replaces by a low-dimensional system of population rate equations (see Methods). Optionally, the mean adaptation current is also recorded at each grid point, as used in the explicit-adaptation example (Supplementary Figure 3). Once *F* is known for each population, the collective dynamics follow from the input rates alone, without reference to individual neurons. It is characterized empirically. A population of uncouples neurons of a single type is driven by independent Poisson inputs, and the excitatory and inhibitory input rates are swept over a grid. At each grid point the stationary output rate *ν*_out_ and its standard deviation *σ*_out_ across the population are recorded (see Methods). Because the neurons are not recurrently connected, each one samples the input-output relationship independently, and the population average estimates *F* directly. The measured surface for each of the three populations is shown in Figure 4 A, D, G (markers).

**Figure 4.**
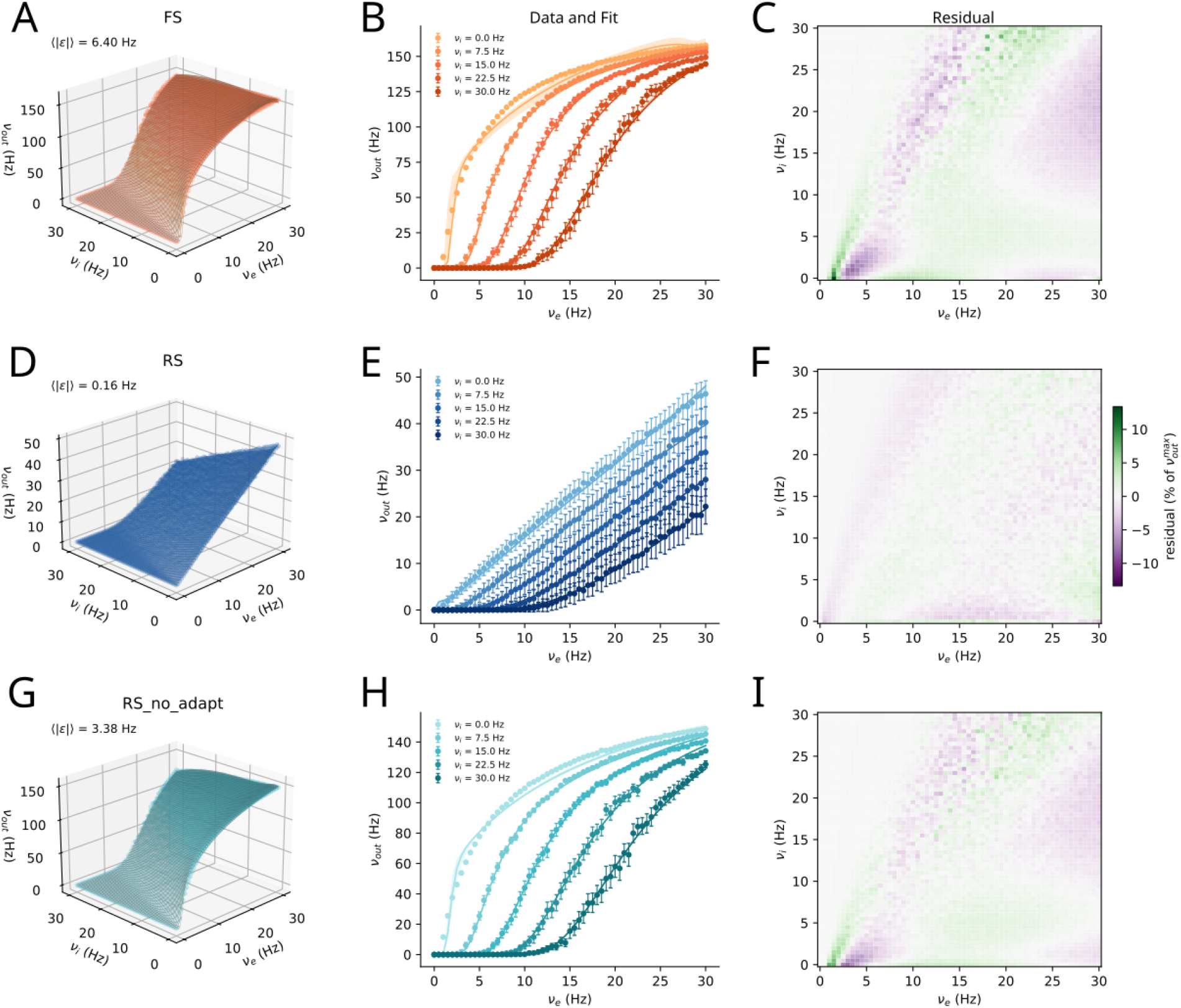
Reconstruction of the transfer function. Rows correspond to the FS (A-C), RS (D-F) and RS_no_adapt (G-I) populations. (A, D, G) Stationary output rate over the grid of excitatory and inhibitory input rates. Markers are simulated output while the surface is the fitted transfer function evaluated on the same grid. The mean absolute error of the fit is reported in each panel. (B, E, H) Same data as function of teh excitatory input rate, at five inhibitory input rates (shades). Markers and error bands represent mean and standard deviation of the simulated output rate, lines are the best fit, and shaded bands are the interval containing the ten retained fits. (C, F, I) Residual, the difference between the simulated and the fitted output rate, expressed as a percentage of the largest simulated output rate of each population. The colour scale is hence common to the three rows.

To obtain a closed form suitable for the population equations, BRIDGE fits the semi-analytical expression of Zerlaut et al. [2018] and Di Volo et al. [2019] to the measured surface. In this form the output rate is written through the mean, standard deviation, and autocorrelation time of the subthreshold membrane potential and a phenomenological effective threshold. The latter expanded as a second-order polynomial whose coefficients are estimated from data (Eqs. 8-10; see Methods for details). The fitted transfer function reproduces the measured surface for all three populations (Figure 4A, D, G, grids), with a mean absolute error reported in each panel. It is important to notice that the procedure does not return a single set of coefficients, but the ten of lowest error, retained for the identifiability analysis.

The one-dimensional slices at fixed inhibitory rate (Figure 4B, E, H) show the fitted curve lying within the dispersion of the simulated rate across the sample range. The band spanned by the retained fits is narrow relative to that dispersion, indicating that the retained parameter sets are not separable by the data. The residual maps (Figure 4C, F, I), expressed as a percentage of the largest simulated rate of each population on a common colour scale, are small over most of the grid and concentrate along the onset front of the transfer function, the diagonal where the output rate rises steeply from silence and where the semi-analytical approximation is least accurate (the residual changes sign across this front, consistent with a small offset in its fitted position). The front is pronounced for FS and RS_no_adapt, which reach output rates above 140 Hz. Adaptation caps the RS output rate near 50 Hz and smooths its input-output surface, so no steep front develops and the residual stays uniformly small ( ⟨|*ϵ*| ⟩= 0.16 Hz).

Because adaptation shapes the RS surface in this way, BRIDGE also supports an alternative characterization in which the adaptation current is included as an explicit input to the transfer function rather than absorbed into the stationary rate (Supplementary Material, Supplementary Figure 3).

### 3.5 Identifiability of the fitted parameters

The fit returns ten sets of parameters corresponding to the lowest error, and a natural question is whether these describe one solution or several distinct ones of comparable quality, and whether the individual coefficients are pinned down by the data. BRIDGE examines this ensemble directly, quantifying how tightly each coefficient is constrained, whether the ten fits fall into separate groups, and along which directions in parameter space they are free to vary (see Methods). The analysis is applied unchanged to the three populations (Figure 5).

**Figure 5.**
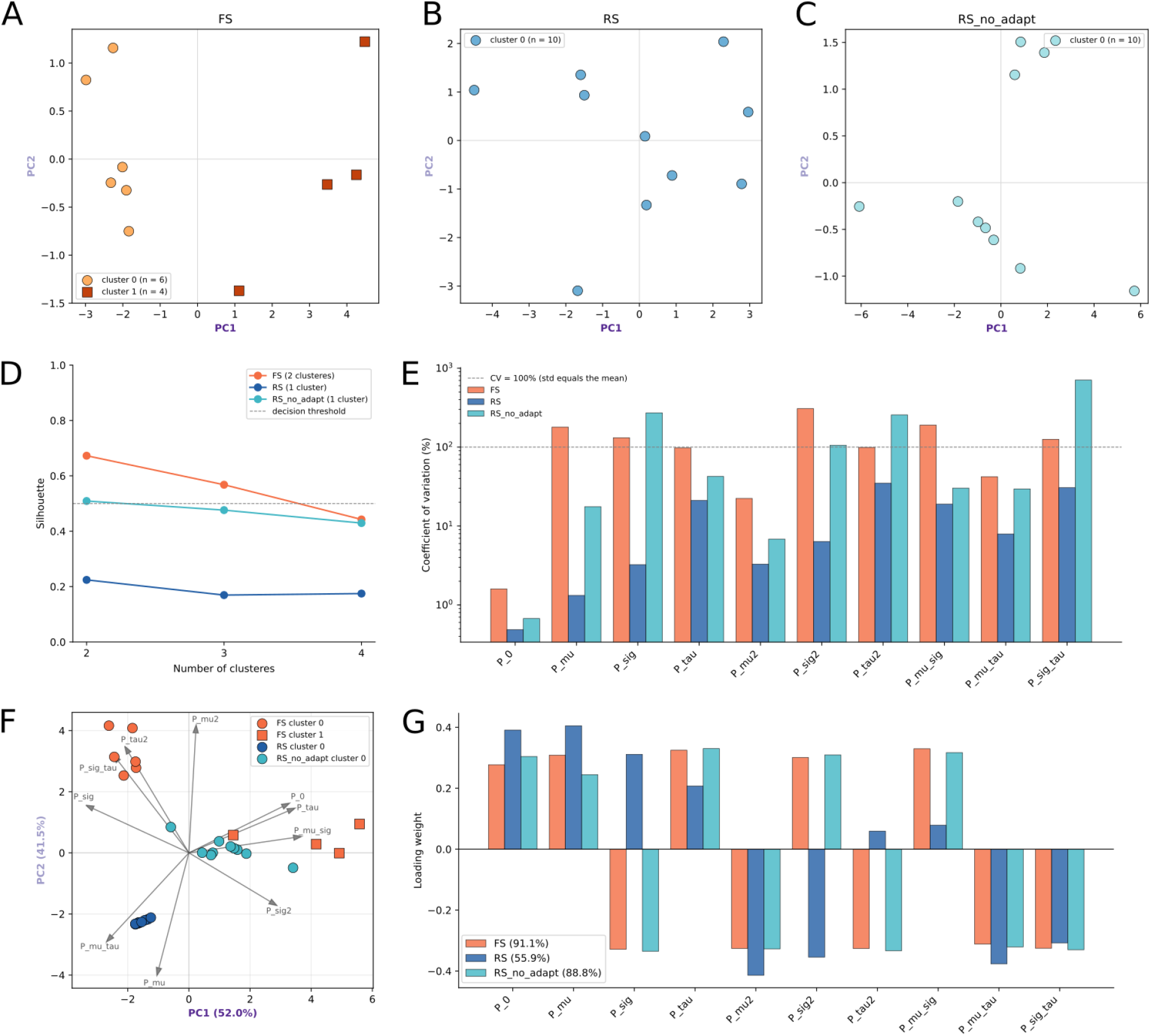
Identifiability of the fitted transfer-function parameters across the three populations. All panels originate from the *N* = 10 lowest error parameter sets retained for each population (FS, orange; RS, blue; RS_no_adapt, steal). A-C) Fits projected onto the first two principal components of each population’s own standardisead coefficients. Markers denote clusters detected by Ward clustering. FS resolves into two clusters, while RS and RS_no_adapt each for a single cluster. D) Silhouette score against candidate cluster count. A cluster is accepted only id the silhouette exceeds 0.55 (dashed line). E) Coefficient of variation of each polynomial coefficient, per population (log scale; dashed lin at CV=100%, where the standard deviation equals the mean). F) All three populations projected onto a single principal component analysis fitted to the pooled, standardized ensembles, with coefficient loadings shown as arrows. FS and RS_no_adapt overlap, whereas RS is displaced and tightly concentrated. G) Loadings of the first principal component, sign oriented to FS. FS and RS_no_adapt share a near-identical component, while RS differs, with sign changes on several coefficients. The first component accounts for 91.1%, 55.9%, and 88.8% for FS, RS and RS_no_adapt, respectively.

The three ensembles are organizes differently (Figure 5A–D). For FS the ten fits split into two separate groups, which represent two distinct solutions of equal quality rather than small perturbations of one. For the RS and RS_no_adapt the fits form a single group. The number of groups is set by clustering score (panel D), and the projection onto each population’s leading directions of variation displays the resulting grouping (panel A-C).

Only one coefficient, the constant term *P*_0_, is well constrained across all three populations, the remaining ones vary by up to two orders of magnitude more while the fitting error stays essentially unchanged (Figure 5E). The individual coefficients are therefore not identifiable, the transfer function is well determined as a surface while the polynomial coefficients that generate it are not. To ask whether the populations share the same unconstrained directions, the three ensembles are compared in a common frame (Figure 5F,G). Projected together, FS and RS_no_adapt overlap, whereas RS is offset and forms a tight, compact group (panel F). The leading direction of variation is near-identical for FS and RS_no_adapt and accounts for most of each population’s spread (91.1% for FS, 88.8% for RS_no_adapt), so their fitting slack lies along a single shared axis (panel G). For RS this direction differs, changing sign on several coefficients, and accounts for only 55.9% of the spread. This means that its slack is distributed across several directions rather than concentrated in one. The compactness of RS in teh common frame is consistent with this. The projection is dominated by the axis shared by the two non-adaptive populations, and RS, whose variation lies largely elsewhere, extends little along it.

### 3.6 The reconstructed mean-field models reproduce the spiking networks within a well-defined regime

BRIDGE validates each reconstructed MF model against its NN by driving the two-population network over a grid of external (excitatory and inhibitory) afferent rates and comparing the stationary population rates predicted by the MF which those measured in the network.

For the FS population the two descriptions agree almost exactly (slope 0.98, RMSE 0.77 Hz over the full grid), the MF matched the network within 15% at nearly every operating point (208 of 212, Figure 6A–D), the fewer larger relative errors being confined to the lowest-drive edge where the rates themselves are small.

**Figure 6.**
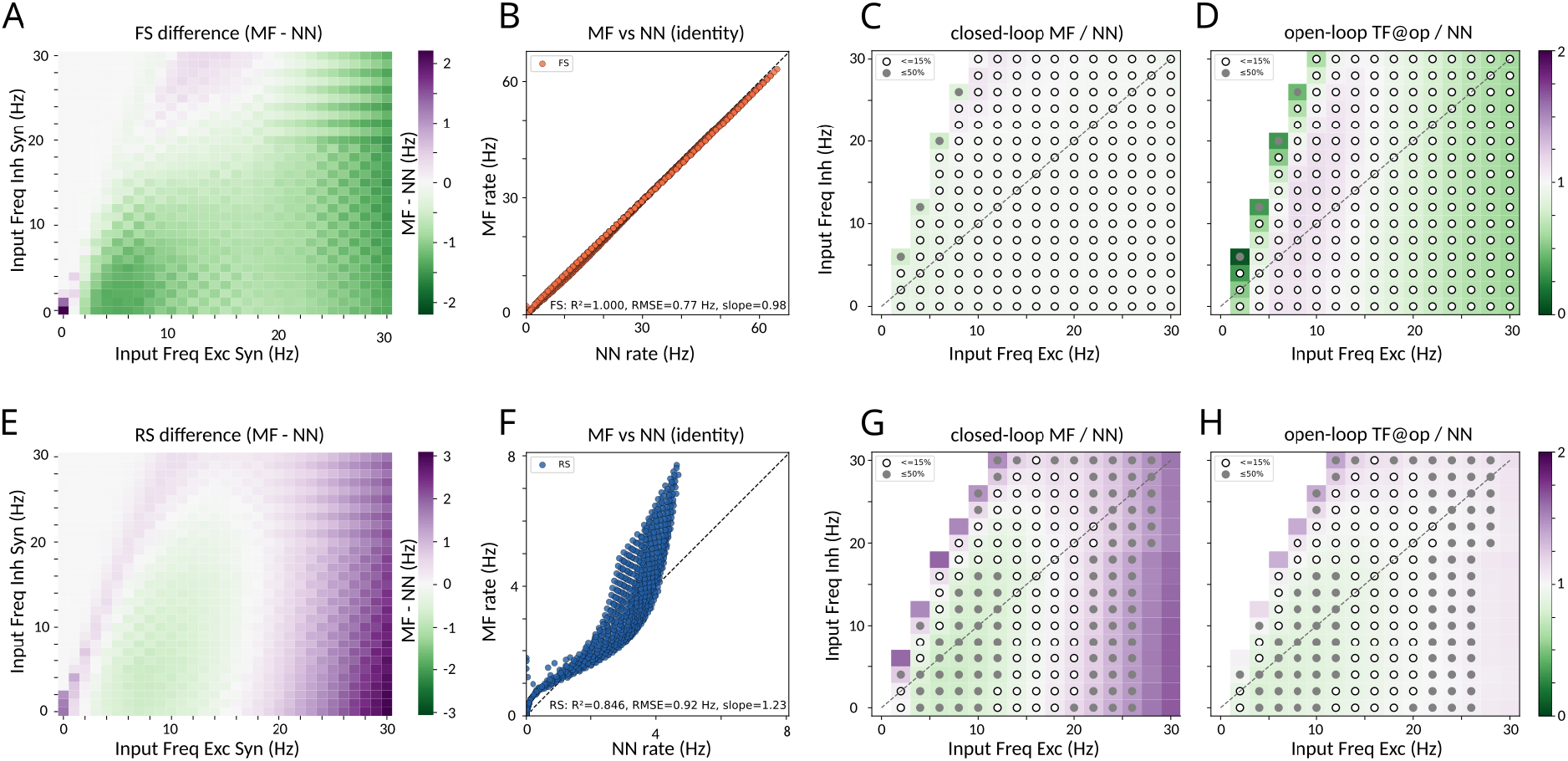
Validation of the reconstructed mean-field models again the spiking networks. Top row (A-D), fast-spiking (FS), bottom row (E-H), regular-spiking (RS) population. All panels span the grid of external excitatory and inhibitory input rates. A, E) difference between the stationary rate predicted by the MF and measured in the NN, *F*_MF_ − *F*_NN_ (Hz); green MF below the network, purple above. B, F) MF versus NN stationary rate at every grid point, and the dashed line represents the identity. Insets contain *R*^2^, RMSE and slope. C, G) closed-loop ratio *F*_MF_*/F*_NN_. D, H) open-loop ratio *F*_TF_*/F*_NN_, where the transfer function evaluated at the network’s measured operating point (external drive plus the recurrent input inferred from the measured rates), which isolates transfer-function accuracy from recurrent amplification. In C, D, G, H, open circles mark quantitative agreement ( ≤ 15%) and fulled circles qualitative agreement ( ≤ 50%). The dashed diagonal marks *ν*_exc_ = *ν*_inh_, and operating points with network rate below 0.5 Hz are omitted. FS agrees quantitatively at nearly every operating point (208*/*212 points within 15%; median 2%; slope 0.98). For RS the transfer function stays close to the network at the operating point (H), while the closed-loop mean-field under-predicts at low-to-moderate excitability and over-predicts as excitatory drive increases (slope 1.23), exceeding 50% error in the high-excitation corner, qualitative agreement (open circles) forms a band at moderate excitation.

For the RS population the MF was in quantitative agreement with the network (relative error ≤ 15%) over a band of moderate excitatory drive (*ν*_exc_ ≈12-20 Hz) that spanned all inhibitory levels (83 of 207 operating points) and in qualitative agreement ( ≤ 50%) held over most of the grid (177 of 207). The open-loop comparison localises the residual. Evaluated directly at the network’s measured operating point, the RS transfer function departed from the network rate far less than the full MF (Figure6, panel H versus G). This means that the transfer function is accurate where the network actually operates. The closed-loop over-prediction at high excitation therefore does not reflect an inaccurate fit but the amplification of a transfer-function residual by the recurrent excitatory loop, and amplification that grows with the effective excitatory gain. The RS non-adaptive population makes this mechanism explicit (Supplementary Figure 4). With spike-frequency adaptation removed (*b* = 0), the closed-loop comparison degraded sharply, and the MF over-predicted the RS rate by a median factor of 15.9, and by up to an order of magnitude, with agreement confined to a few near-balanced operating points. Removing adaptation raises the effective recurrent excitatory gain, and it is in this high-gain regime that the MF description, together with its amplification of the transfer function residuals, is least accurate.

Across the three populations, MF accuracy is therefore set by the feedback that opposes recurrent excitatory rather than by the quality of the fit. It is highest for the inhibitory population, whose recurrence is intrinsically self-stabilizing, it remains quantitative for the adaptive RS population over a defined range of moderate drive, where adaptation supplies a negative feedback that limits the effective gain, and it deteriorates for the non-adaptive excitatory population, in which recurrent excitation is unopposed. The MF is the most predictive in the inhibitory or adaptation-stabilized fluctuation-driven regime, and least predictive in the excitation-dominated regime where the recurrent gain is highest.

BRIDGE allows a user, reconstructing a mean-field model from their own recordings, to delineate the regime of validity and attribute any shortfall either to the transfer function fit or to the recurrent dynamics.

### 3.7 Stationary dynamics and stability of the mean-field network

To investigate the collective behavior of the reconstructed MF, BRIDGE also characterizes its stationary dynamics through a phase-plane representation (Figure 7). Unlike the conventional inspection of firing-rate time series, which reports how each population rate evolves as a function of time, the phase plane represents the instantaneous state of the coupled system as a point in state space, here defined by (*ν*_FS_, *ν*_RS_), thereby making the geometry of the underlying dynamical system explicit. This representation makes it possible to distinguish transient motion from stationary behavior, identify the states toward which trajectories converge, and relate these behaviors directly to the nullclines and their intersections: the *ν*_FS_- and *ν*_RS_-nullclines define the loci at which the corresponding population rates are stationary, so that their intersections correspond to fixed points of the coupled mean-field dynamics, revealing how recurrent excitation and inhibition jointly shape the trajectory through population-rate space.

**Figure 7.**
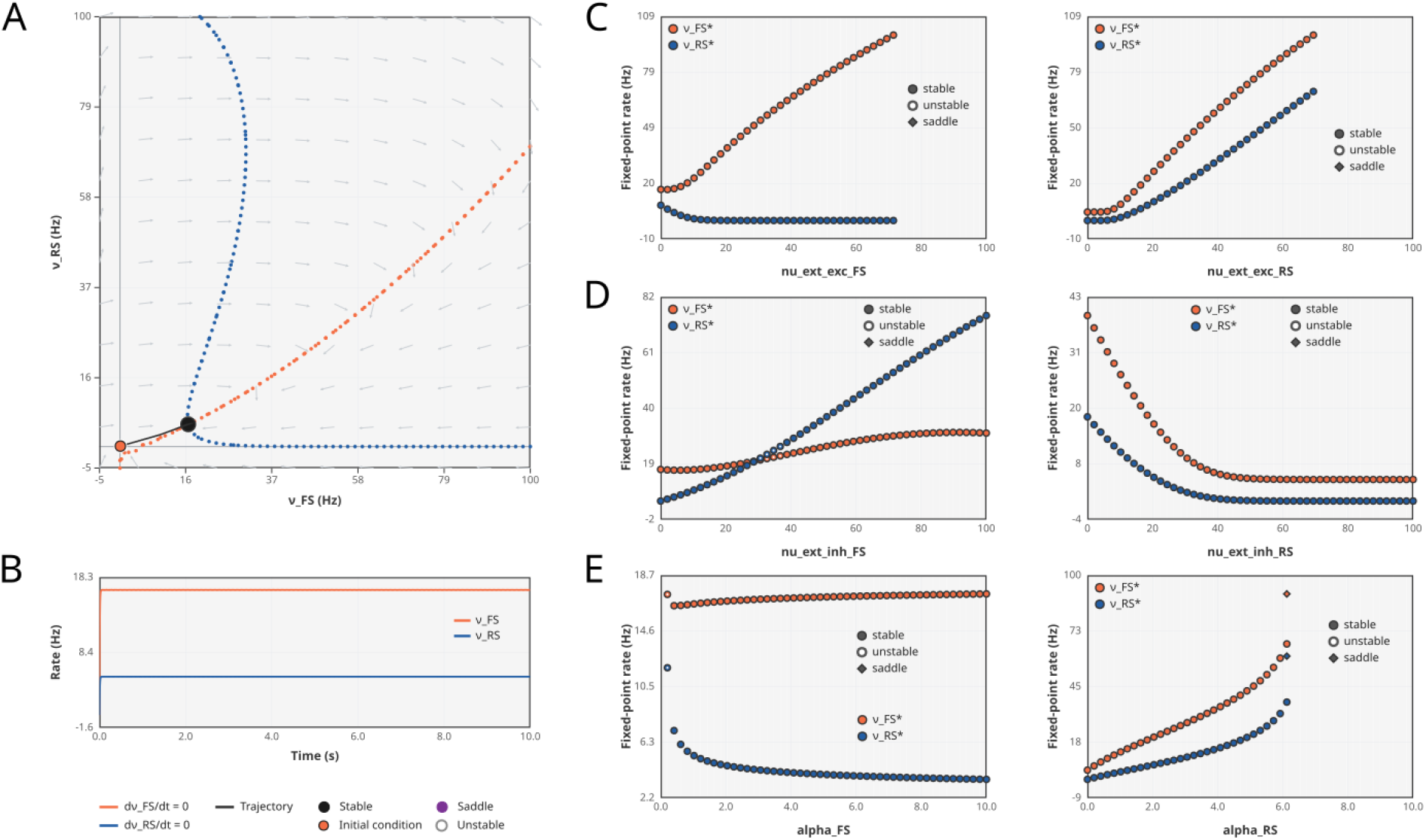
Dynamical characterization of the mean-field network. Phase plane of the two-population mean-field model in the (*ν*_FS_, *ν*_RS_) plane. The FS nullcline (d*ν*_FS_*/*d*t* = 0, orange) and the RS nullcline (d*ν*_RS_*/*d*t* = 0, blue) intersect at the network equilibria, each classified by linear stability as a stable node, an unstable node, or a saddle. A representative trajectory, integrated from the marked initial condition, relaxes onto a stable fixed point. B) Temporal realization of the trajectory in A. The population rates *ν*_FS_ (orange) and *ν*_RS_ (blue) evolve from the initial transient to their stationary values. C-E) One-parameter continuation of the network equilibria. Each panel plots the fixed-point rates 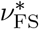 (orange) and 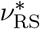 (blue) against a control parameter applied to the FS (left column) or RS (right column) population, with branch stability coded as stable, unstable, or saddle. C) External excitatory afferent rate, D) external inhibitory afferent rate, E) *α* scaling factor.

Under a baseline drive the two nullclines intersect at a single low-rate equilibrium in the (*ν*_FS_, *ν*_RS_) plane, and a representative trajectory initiated away from this equilibrium follows the local flow toward the intersection, converging onto this stable fixed point (Figure 7A). Its temporal profile shows a brief onset transient followed by relaxation to the stationary rates, with FS settling above RS (Figure 7B), demonstrating that this transient is not simply a sequence of independent rate changes, but the relaxation of the coupled system toward an attracting state.

We next trace how this equilibrium reorganizes under sustained changes in afferent drive by continuing the fixed points along each external input in turn (Figure 7C–E). Raising the excitatory afferent rate to FS increased 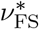 while holding the RS near quiescence, whereas the same drive applied to RS raised both rates, with the excitatory population pulling FS upward through recurrent excitation onto it. Inhibitory drive produced the complementary picture: inhibiting FS disinhibited RS, so 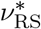 rose and 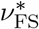 fell, while inhibiting RS suppressed both populations together.

Over most of these ranges the equilibrium is stable, however the continuation also resolved localized changes of stability. The equilibrium lost stability near an external inhibitory rate to FS of ≈ 38 Hz, where both branches became unstable (Figure 7D, left). The *α* continuation showed the same structure: both branches were unstable near *α*_FS_ → 0 (Figure 7E, left), and saddle equilibria emerged on both branches at *α*_RS_ ≈ 6 (Figure 7E, right). These instabilities delimit the parameter regime over which the MF model sustains a single attractor.

More importantly, the phase-plane formulation provides a natural framework for examining how the qualitative organization of the dynamics changes with model parameters, beyond the specific quantitative trends reported above. Rather than treating each continuation as an isolated numerical result, this framework transforms a collection of rate trajectories into a mechanistic description of the state-space geometry governing those trajectories, providing direct access to attractors, stability boundaries, and the parameter regimes in which qualitatively different collective dynamics can emerge.

### 3.8 Networks of coupled mean-field models

The validated mean-field models can be assembles into networks. BRIDGE allows the construction of networks of arbitrarily many populations and nodes. It just requires the network configuration: each population needs to be assigned a type (with its single cell and fitted transfer function parameters), a node, and each directed connection with given probability, receptor type, synaptic strength, and transmission delay.

A two-node example is shown in Figure 8. Node 1 contains three populations comprising FS (FS1), RS (RS1), and RS_no_adapt (RS2), while Node 2 is a two-population module with FS (FS2) and RS (RS3). The nodes are couples by a single long-range excitatory projection from RS1 to RS3 with a 10ms conduction delay. All populations receive a common fluctuating background drive (Ornstein-Uhlenbeck noise), and under this drive both nodes settle into a low-rate baseline (Figure 8A). To investigate signal transmission, a targeted input is added on top of the background activity and applied to one neuronal population at a time. Three temporal profiles are shown: a Gaussian transient into RS1 (Figure 8B), an oscillatory drive (sinusoid Ornstein-Uhlenbeck signal) into RS2 (Figure 8C), and a purely stochastic Ornstein-Uhlenbeck drive into FS1 (Figure 8D). In each case the input recruits the driven population and propagates through the local recurrent connections to the remainder of Node 1 and, via the long-range projection, to Node 2, where it arrives attenuated. The downstream effect depends on the identity of the driven population: exciting RS1 and RS2 raises the excitatory output of Node 1 and thereby the activity of Node 2, while driving the inhibitory FS1 suppresses the local excitatory rates and, in turn, lowers the activity of Node 2.

**Figure 8.**
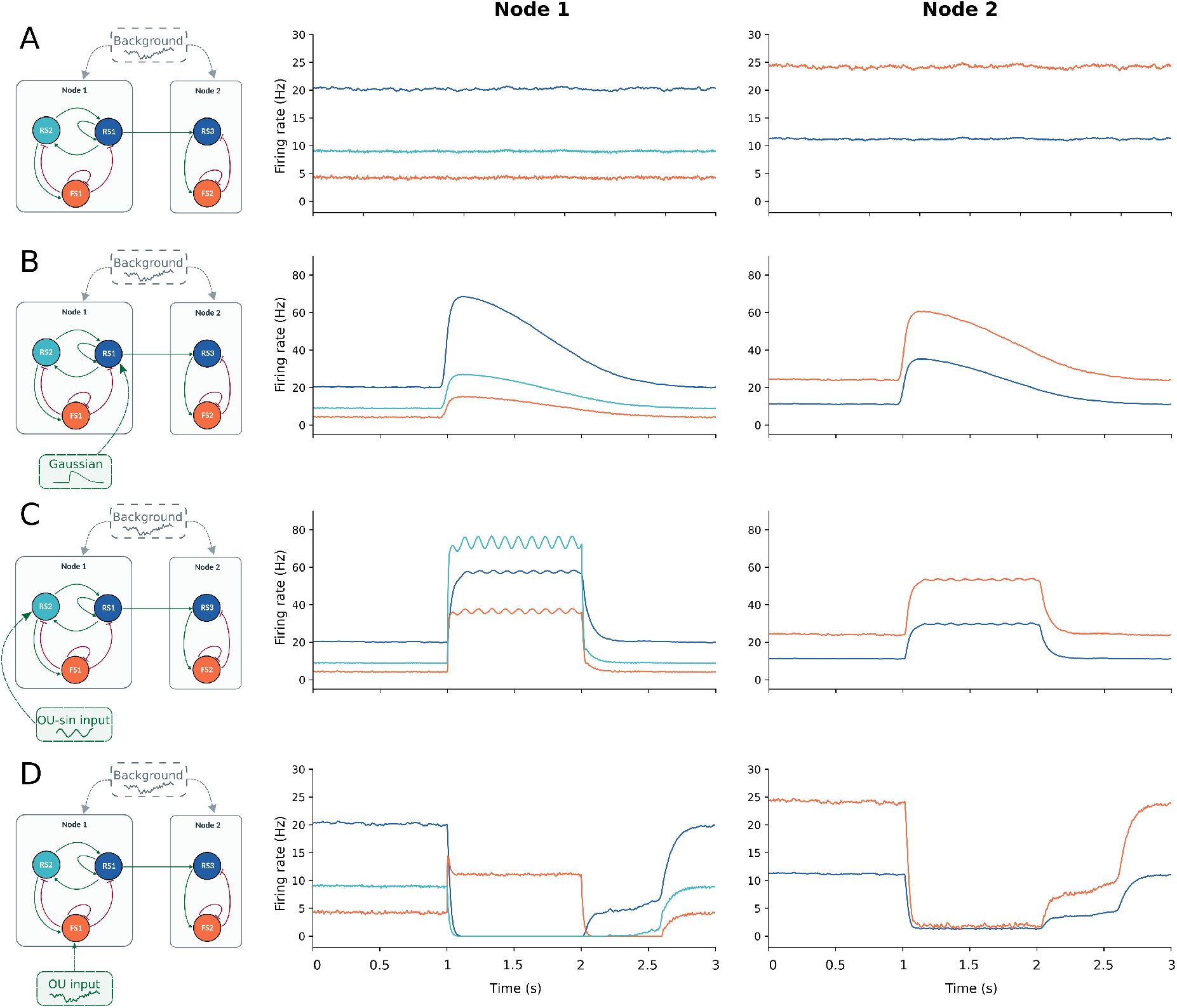
Network of mean-field models. Each row corresponds to a different external drive; each columns shows, from left to right, the network topology (with driven population indicated by the external-input arrow), the population firing rates of Node 1, and the population firing rates of Node 2. Node 1 contains an adaptive excitatory population RS_1_ (blue), a fast-spiking inhibitory population FS_1_ (orange), and a non-adaptive excitatory population RS_2_ (teal). Node 2 contains an adaptive excitatory population RS_3_ (blue) and an inhibitory population FS_2_ (orange), The nodes are coupled by a single feedforward excitatory projection RS_1_ → RS_3_ with a 10 ms conduction delay, and all population receive a common Ornstein-Uhlenbeck (OU) background drive. A) Background drive only, giving a low-rate baseline. B-D) Targeted signal is added into a single population, a Gaussian transient into RS_1_, and oscillatory (sinusoidal OU) drive into RS_2_, and a OU drive into FS_1_ respectively. The full network configuration (nodes, populations, connection probabilities, receptor types, delays) is defined in a single JSON file which can be edited and passed to the corresponding notebook to define a new network (for example containing more nodes) and study signal propagation.

## 4 Discussion

We introduce BRIDGE, a modular and open-source workflow for reconstructing, analyzing, validating, and coupling mean-field models starting from single-neuron dynamics. Its principal contribution is not a new mean-field formalism in itself, but the integration of steps that are commonly implemented independently, together with built-in analytical tools that allow users to inspect and evaluate each stage of the derivation. By fixing this derivation path while leaving the neuronal model, synaptic channels, connectivity, and population composition configurable, BRIDGE turns a case-specific modeling procedure into a reproducible and reusable workflow.

Mean-field derivations have traditionally required substantial analytical and numerical work for each new cellular model or circuit. Even when based on the same theoretical formalism, implementations often differ in their conventions, data structures, fitting procedures, and validation criteria, making models difficult to reproduce or transfer between studies. BRIDGE addresses this methodological fragmentation by retaining the intermediate products of the derivation and exposing them to inspection. The user therefore obtains not only a final system of population-rate equations but also the intermediate stages, each accessible through built-in analysis tools.

We illustrate these capabilities using three neuronal populations (FS, RS and RS_no_adapt), not to report properties of these particular models, but to demonstrate the kind of characterization BRIDGE makes routine.

One example is the distinction between identifiability of the transfer-function surface and identifiability of its polynomial coefficients. Across the three populations, several substantially different coefficient sets produced nearly indistinguishable input–output surfaces and comparable fitting errors. The transfer function can be sufficiently constrained for population simulations even when its individual phenomenological coefficients are not uniquely determined. This distinction would be hidden by reporting only the single parameter set with the lowest error.

The retained ensembles nevertheless contained informative structure. The FS fits separated into two groups of comparable quality, while the RS and RS_no_adapt fits each formed a single group. Moreover, the two non-adaptive populations shared a dominant direction of parameter variation, whereas the adaptive RS population exhibited a different and more distributed structure.

The validation procedure provides a second important diagnostic contribution. A discrepancy between a mean-field model and its spiking-network counterpart can arise because the fitted transfer function is inaccurate at the relevant input, because recurrent interactions amplify a small local residual, or through a combination of both mechanisms. BRIDGE separates these possibilities by combining open-loop and closed-loop comparisons. The open-loop evaluation tests the transfer function at the operating point measured directly in the spiking network, whereas the closed-loop evaluation tests the complete self-consistent mean-field system. Their comparison therefore identifies whether improvement should focus on refitting the single-population closure or on characterizing the recurrent regime in which that closure is embedded.

This distinction was particularly evident for the adaptive RS population. The transfer function remained relatively accurate when evaluated at the input received by the spiking network, while the closed-loop mean-field increasingly over-predicted the population rate at high excitatory drive. The disagreement therefore resulted primarily from recurrent amplification of a comparatively small transfer-function residual rather than from a uniformly poor fit. The non-adaptive RS network provided a more extreme example of the same mechanism: removing spike-triggered adaptation increased the effective excitatory gain and strongly degraded closed-loop agreement. In contrast, the inhibitory FS population remained accurately reproduced over nearly the complete input domain examined. These comparisons show that the accuracy of a reduced population model cannot be inferred from transfer-function fitting error alone. It also depends on how the local error is transformed by the feedback architecture of the recurrent network. BRIDGE describes the accuracy of a mean-field model through a *regime of validity*, reporting quantitative and qualitative agreement, measures that are rarely reported.

Beyond reproduction of stationary firing rates, BRIDGE provides access to the geometrical organization of the reduced dynamics. The phase-plane analysis identifies nullclines, fixed points, their stability, and trajectories through population-rate space, while parameter continuation tracks how this organization changes with afferent input or model parameters. More generally, these tools can be used to identify multistability, transitions between activity states, changes in excitability, or parameter regions associated with pathological dynamics. The reduction therefore provides not only computational efficiency but also a dynamical-systems representation through which network behaviour can be interpreted mechanistically.

The final network layer extends this approach from an isolated local circuit to coupled meanfield nodes. Importantly, nodes in BRIDGE are not required to contain identical populations or share the same transfer functions. Population composition, neuronal parameters, receptor channels, connection weights, and delays can be specified independently, making it possible to construct large-scale models in which local dynamics differ between brain regions.

We demonstrated BRIDGE primarily using the AdEx model, chosen for its balance between dynamical richness, computational efficiency, and biological interpretability. The modular architecture does not depend on this choice: alternative single-neuron models can be substituted by supplying their equations and reset conditions together with the corresponding synaptic channels, so that deriving a mean-field model for a new cell type reduces to characterizing its transfer function within the same workflow rather than re-implementing the derivation from scratch. We exercised this generality on the E-GLIF model of cerebellar neurons (Lorenzi et al. [2023]). The value of such reproducible derivations is reflected in concurrent efforts to automate the construction of mean-field models: Lorenzi et al. [2026] recently proposed an automated derivation of mean-field models from spiking networks simulated in NEST. BRIDGE shares this goal of turning a case-specific derivation into a systematic procedure, and complements it through its Python/Brian2 implementation and its emphasis on built-in analysis, identifiability, open- and closed-loop validation, and phase-plane characterization, together with the extension to networks of heterogeneous mean-field nodes.

The current implementation of BRIDGE inherits several limitations from the underlying mean-field formalism. The semi-analytical closure assumes asynchronous irregular, fluctuation-driven activity, and its accuracy degrades in the mean-driven regime and along the low-inhibition edge of the input grid where this assumption is weakest. The reduction demonstrated here is first-order, tracking only the mean population rates; a second-order description that also propagates the rate covariances would extend the analysis to finite-size fluctuations.

Despite these limitations, BRIDGE establishes a practical route across modelling scales. It begins with microscopic neuronal and synaptic descriptions, constructs and validates a mesoscopic population representation, and then enables those population models to be embedded in large-scale networks. Its broader value lies not only in making mean-field models easier to construct, but in making their derivation inspectable, their limitations measurable, and their use in large-scale simulations more transparent. Finally, to support adoption and reproducibility, BRIDGE will be made available through the EBRAINS research infrastructure.

## 5 Code availability

The implemented codes and detailed descriptions are available here: https://github.com/IlaCar/neuron-to-meanfield.

Simulations were performed on a Linux-based desktop computer equipped with an AMD Ryzen 9 3900X 12-core (24-thread) processor and 64 GB of RAM.

## 6 Acknowledgments

IC thanks Dr Roberta Maria Lorenzi and Dr Alice Geminiani for introducing her to the fascinating field of cerebellar modelling and for sharing their expertise and the information required to extend BRIDGE to support the complete E-GLIF modelling workflow.

IC: Digital Futures. AD: Research supported by CNRS, Agence Nationale de la Recherche (FLAG-ERA BrainAct project, CR-CNS ImpactCom project), and the European Union (Human Brain Project H2020-945539 and Virtual Brain Twin project 101137289). DD: The preparation of this article was funded through the EU’s Horizon Europe Programme SGA 101147319 (EBRAINS 2.0), SGA 101137289 (Virtual Brain Twin), and No. 101057429 (project environMENTAL), and government grant managed by the Agence Nationale de la Recherche reference ANR-22-PESN-0012 (France 2030 program). PP: Supported by the Virtual Brain Twin project, funded by the European Union under Horizon Europe Grant Agreement No. 101137289.

## 7 Author contributions

IC, and AD conceived the study. IC built the pipeline developing the methodology. IC performed the simulations, carried out the investigation, analysis, and prepared the visualizations. DD initiated the phase plane analysis. MW reformatted the GitHub folder, provided initial code for phase plane widget. PP developed the network of mean-field construction and provided the initial code. IC wrote the original manuscript draft. DD, MW, PP, and AD contributed to review and editing.

