## Supplementary Material for "BRIDGE: A Computational Workflow from Single Neurons to Network of Mean-Field Models"

#### 1 E-GLIF neuron model and alpha-synapses

BRIDGE is not tied to the AdEx neuron with conductance-based synapses used in the main text. To demonstrate this, we reconstruct a mean-field model from a different single-neuron formalism, the E-GLIF model Geminiani et al. [2018], combined with alpha synapses.

The E-GLIF framework describes the evolution of the membrane potential $V_m(t)$ of a single-compartment neuron together with two intrinsic currents: an adaptive current $I_a(t)$ and a depolarizing spike-triggered current $I_d(t)$. It consists of a system of three differential equations:

$$\begin{cases} \dfrac{dV_m}{dt} = \dfrac{1}{C_m}\left(-\dfrac{C_m}{\tau_m}(V_m(t) - E_L) - I_a(t) + I_d(t) + I_e + I_{stim} - I_{syn}(t)\right) \\ \dfrac{dI_a}{dt} = k_a(V_m(t) - E_L) - k_2\, I_a(t) \\ \dfrac{dI_d}{dt} = -k_1\, I_d(t) \end{cases} \tag{1}$$

where $C_m$ is the membrane capacitance, $\tau_m$ the membrane time constant, $E_L$ the resting potential, $I_e$ a constant endogenous current modelling the net contribution of depolarizing ionic currents that generate autorhythmicity, $I_{stim}$ the external (injected) stimulation current, $I_{syn}(t)$ the total synaptic current, $k_a$ and $k_2$ the adaptation constants, and $k_1$ the decay rate of the depolarizing current.

If the neuron is not in the refractory interval $t_{ref}$, a spike is generated stochastically at $t_{spk}$ by a point process with escape rate $\lambda(t)$:

$$\lambda(t) = \lambda_0 \exp\left(\frac{V_m(t) - V_{th}}{\tau_V}\right), \tag{2}$$

where $V_{th}$ is the threshold potential and $\lambda_0$, $\tau_V$ are the escape-rate parameters. Spike initiation is therefore stochastic rather than a hard threshold crossing: $\lambda_0$ sets the instantaneous spike rate when $V_m = V_{th}$, and $\tau_V$ sets the voltage scale over which the rate varies, so that smaller $\tau_V$ approaches a deterministic threshold.

When a spike occurs at $t_{spk}$, the state variables are reset according to:

$$\begin{cases} V_m \to V_{reset} \\ I_a \to I_a + A_2 \\ I_d \to A_1 \end{cases} \tag{3}$$

where $V_{reset}$ is the reset potential and $A_1$, $A_2$ are the current-update constants.

A neuron can be driven by current injection ($I_{stim}$) and/or by synaptic activation. Here we use alpha synapses, whose conductance following a presynaptic spike at $t_{spk}$ is

$$g_{syn}(t) = G_{syn}\,\frac{t - t_{spk}}{\tau_{syn}}\,\exp\!\left(1 - \frac{t - t_{spk}}{\tau_{syn}}\right), \tag{4}$$

with peak conductance $G_{syn}$ and time-to-peak $\tau_{syn}$. Excitatory and inhibitory synapses differ through their reversal potential and time constant. The total synaptic current entering Eq. (1) is the sum over all afferent synapses,

$$I_{syn}(t) = \sum_s g_{syn}^{(s)}(t)\,\big(V_m(t) - E_{syn}^{(s)}\big), \qquad E_{syn}^{(s)} \in \{E_{ex}, E_{in}\}, \tag{5}$$

where $E_{ex}$ and $E_{in}$ are the excitatory and inhibitory reversal potentials. With the negative sign of $I_{syn}$ in Eq. (1), excitatory input ($E_{ex} > V_m$) depolarizes the membrane and inhibitory input hyperpolarizes it.

The same construction extends to an arbitrary number of afferent populations: each presynaptic population $s$ contributes its own alpha conductance with population-specific parameters ($G_{syn}^{(s)}$, $\tau_{syn}^{(s)}$) and reversal potential $E_{syn}^{(s)}$, and the total synaptic current is the sum of these contributions,

$$I_{syn}(t) = \sum_s g_{syn}^{(s)}(t)\,\big(V_m(t) - E_{syn}^{(s)}\big). \tag{6}$$

Here we apply this to cerebellar Golgi cells, which receive three afferent populations: two excitatory (granule-cell and mossy-fiber) and one recurrent inhibitory (Golgi–Golgi) input, giving a three-dimensional transfer function.

#### Mean-field reduction

Following the same semi-analytical procedure used in the main text (Zerlaut et al. [2018], Di Volo et al. [2019]), the mean-field description of a population of E-GLIF neurons with alpha synapses was derived by Lorenzi et al. [2023]; here we adopt their formulation within BRIDGE.

As before, the population output rate is set by a transfer function $F$ that maps the mean afferent rates $\nu_s$ of the presynaptic populations onto the stationary firing rate. $F$ retains the fluctuation-driven, erfc-based form; the alpha synaptic kernel enters only through the membrane-potential moments. For the Golgi cell this generates a three-dimensional transfer function, with two excitatory afferents (granule-cell and mossy-fiber) and one recurrent inhibitory afferent.

The mean input conductance and the effective membrane time constant are

$$\mu_G = g_L + \sum_s K_s\,Q_s\,\tau_s\,\nu_s, \qquad \tau_m^{\text{eff}} = \frac{C_m}{\mu_G}, \tag{7}$$

where, for each presynaptic population $s$, $K_s$, $Q_s$ and $\tau_s$ are the synaptic convergence, quantal conductance and decay time. The first three moments of the membrane-potential fluctuations are

$$\mu_V = \frac{1}{\mu_G}\left(e\sum_s K_s Q_s \tau_s \nu_s\,E_s + g_L E_L - W\right), \tag{8}$$

$$\sigma_V^2 = \sum_s \left(2\tau_m^{\text{eff}} + \tau_s\right)\left(\frac{e\,U_s\,\tau_s}{2\left(\tau_s + \tau_m^{\text{eff}}\right)}\right)^2 K_s\,\nu_s, \tag{9}$$

$$\tau_V = \frac{1}{2\,\sigma_V^2}\sum_s K_s\,\nu_s\,\left(e\,U_s\,\tau_s\right)^2, \qquad U_s = \frac{Q_s}{\mu_G}\,(E_s - \mu_V), \tag{10}$$

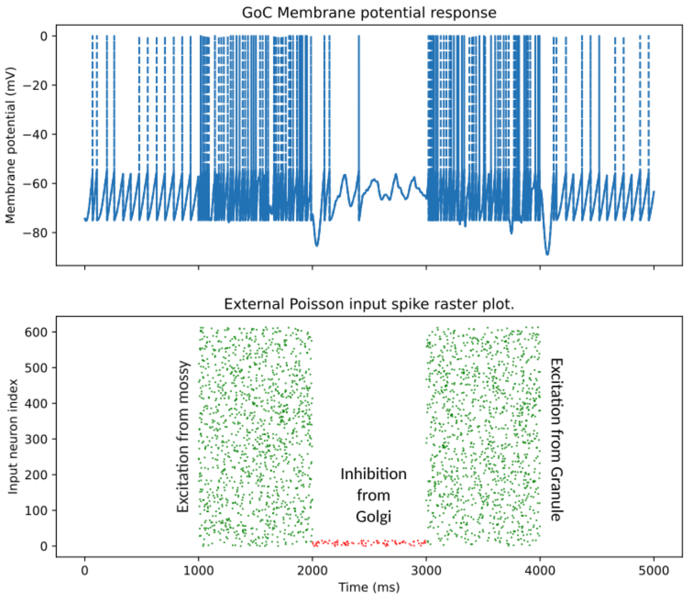


Figure 1: **E-GLIF Golgi cell response to synaptic stimulation.** Membrane-potential response of a Golgi neuron (GoC) modelled using the extended generalized leaky integrate-and-fire (E-GLIF) formalism with alpha-synapses. The upper panel shows the GoC membrane potential, while the lower the raster plot of the external Poisson spike trains delivered through the three synaptic pathways. Excitatory mossy-fiber input is applied from 1 to 2 s, recurrent inhibitory input from 2 to 3 s, and excitatory input from granule cells from 3 to 4.

where $E_s$ is the reversal potential of afferent $s$, $W$ the (stationary) adaptation current, and the factor $e = \exp(1)$ arises from the peak-normalised alpha kernel, replacing the exponential-synapse expressions of the main text.

The transfer function is then

$$F(\nu_s) = \frac{1}{2\,\tau_V^N}\,\frac{g_L}{C_m}\,\mathrm{erfc}\!\left(\frac{V_{\mathrm{thre}}^{\mathrm{eff}} - \mu_V}{\sqrt{2}\,\sigma_V}\right)\alpha, \qquad \tau_V^N = \tau_V\,\frac{g_L}{C_m}, \tag{11}$$

with the effective threshold expressed, as in the main text, as a first-order polynomial in the fluctuation moments,

$$V_{\mathrm{thre}}^{\mathrm{eff}} = P_0 + P_1\frac{\mu_V - \mu_V^0}{\delta\mu_V} + P_2\frac{\sigma_V - \sigma_V^0}{\delta\sigma_V} + P_3\frac{\tau_V^N - \tau_V^{N,0}}{\delta\tau_V^N} + P_4\,\ln\frac{\mu_G}{g_L}, \tag{12}$$

using the normalisation constants of Zerlaut et al. [2018]. The coefficients $P_0, \ldots, P_4$ are fitted to the numerical transfer function following the same two-stage procedure as in the main text, and $\alpha$ is optimised per population to extend the fit to high firing rates.

Coupling these transfer functions preserves the second-order Markovian formalism (El Boustani and Destexhe [2009], Di Volo et al. [2019]) unchanged:

$$\begin{aligned} T\,\frac{d\nu_\mu}{dt} &= (F_\mu - \nu_\mu) + \tfrac{1}{2}\,c_{\lambda\eta}\,\frac{\partial^2 F_\mu}{\partial\nu_\lambda \partial\nu_\eta}, \\ T\,\frac{dc_{\lambda\eta}}{dt} &= \delta_{\lambda\eta}\,\frac{F_\lambda\,(1/T - F_\lambda)}{N_\lambda} + (F_\lambda - \nu_\lambda)(F_\eta - \nu_\eta) + \frac{\partial F_\lambda}{\partial\nu_\mu}\,c_{\eta\mu} + \frac{\partial F_\eta}{\partial\nu_\mu}\,c_{\lambda\mu} - 2\,c_{\lambda\eta}, \end{aligned} \tag{13}$$

so that only the transfer function itself changes when the single-neuron and synaptic models are replaced.

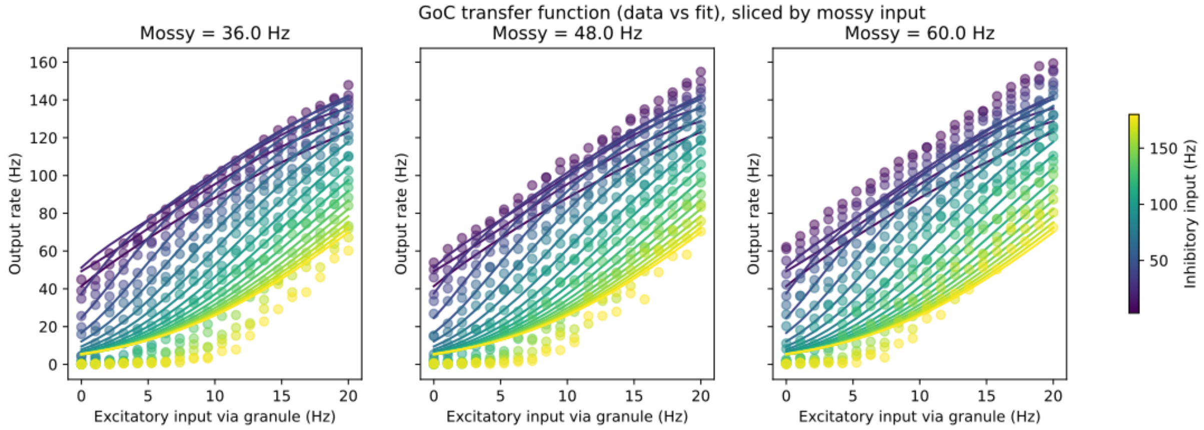


Figure 2: **Transfer function reconstruction.** The transfer function of the GoC neurons is three-dimensional since they receive three distinct input pathways. Here, it is displayed as two=dimensional slices at fixed mossy-fiber input rates of $36, 48$, and 60 Hz. Within each panel, the granule-cell excitatory input rate is shown on the x-axis, the recurrent inhibitory input rate is encoded bu color, and the resulting output firing rate is shown on the y-axis. Markers represent the firing rates obtained from numerical simulations, whereas solid lines show the corresponding fitted transfer function.

#### 2 Transfer function characterization and fitting of the adaptive regular-spiking (RS) population with explicit adaptation.

In the main text the RS transfer function is measured on the stationary firing rate: the first second of each simulation is discarded to remove the adaptation transient, so that adaptation enters the fit only implicitly, through its effect on the stationary output rate. Here we instead record the mean adaptation current explicitly at each grid point and fit it as an additional observable; because adaptation is tracked directly rather than allowed to settle, the entire simulation interval is retained and no transient is discarded. This provides an alternative characterization of the same population and illustrates that the adaptation current can itself be a measured output of the workflow.

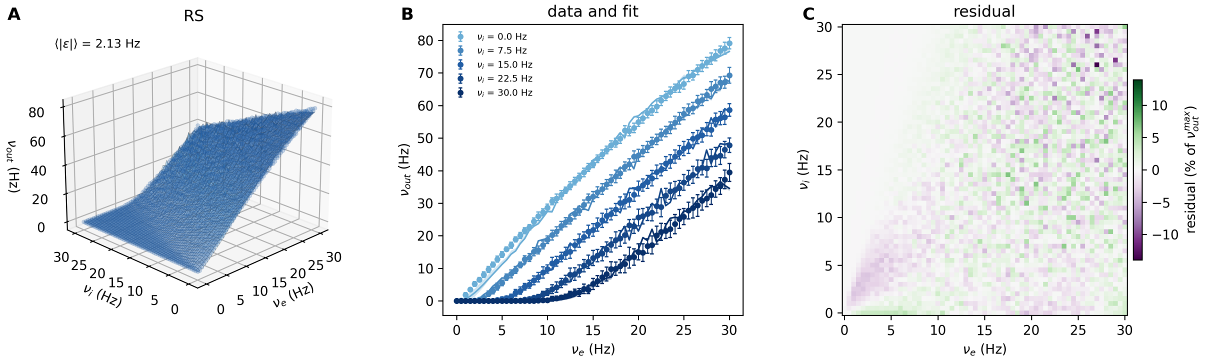


Figure 3: **Transfer function characterization and fitting.** A) Output rate $\nu_{\text{out}} = F(\nu_e, \nu_i, w)$, averaged over the full simulation interval, as a function of the excitatory and inhibitory presynaptic rates; the adaptation current $w$ measured at each grid point is the third input to $F$. Markers are the measured surface; the mesh is the fitted transfer function. B) One-dimensional slices of the same data at fixed inhibitory rates $\nu_i \in \{0, 7.5, 15, 22.5, 30\}$ Hz (markers, measured; lines, fit), with mean absolute error $\langle|\epsilon|\rangle = 2.13$ Hz. C) Fit residual across the $(\nu_e, \nu_i)$ grid, expressed as a percentage of the maximum measured output rate $\nu_{\text{out}}^{\text{max}}$.

### 3 Regime of validity of the RS with adaptation removed (b=0)

The reconstructed MF model for the non-adaptive RS population is validated against its spiking network over the grid of external excitatory and inhibitory afferent rates, as in the main validation Figure 6. Each marker is one operating point; filled markers denote quantitative agreement (relative error $\leq 15\%$) and open markers qualitative agreement ($\leq 50\%$).

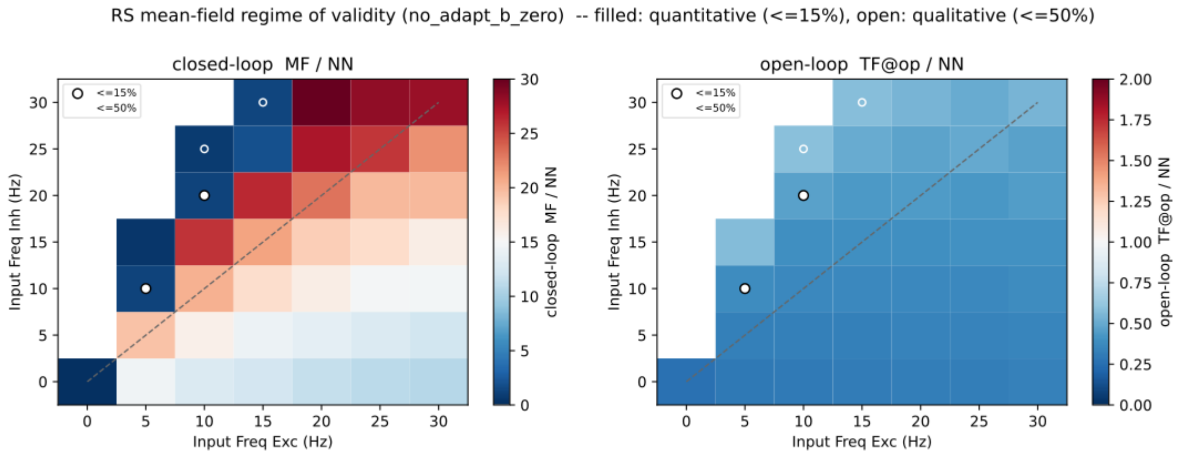


Figure 4: **Transfer function characterization and fitting.** Left) Closed-loop comparison: ratio of the stationary rate predicted by the full mean-field model to the network rate (MF / NN). Right) Open-loop comparison: ratio of the transfer function evaluated at the network's measured operating point to the network rate (TF@op / NN).
The two colour scales differ: the closed-loop scale extends to 30 (saturating there) to accommodate the strong over-prediction, whereas the open-loop scale spans 0–2, since the transfer function at the operating point stays close to the network rate. With adaptation removed the closed-loop model over-predicts across almost the entire grid, while the open-loop transfer function remains near the network.

### 4 Network of 3 mean-fields

A coupled network of three mean-field nodes (N1, N2, N3) reconstructed with BRIDGE, illustrating that the workflow extends to more than two nodes and to nodes of heterogeneous composition.

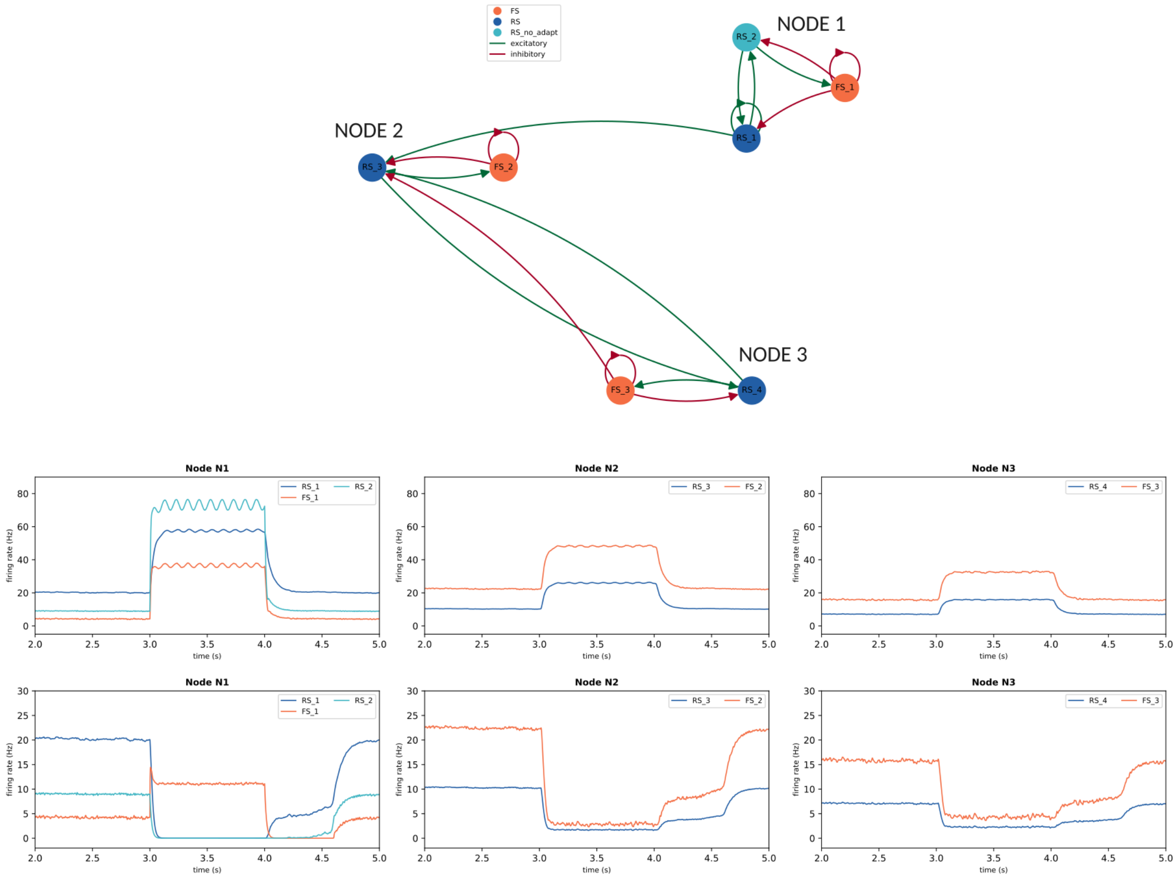


Figure 5: **Signal propagation in a three-node network of mean-field models.** The schematic shows the network topology: node N1 comprises the populations $RS_1$, $RS_2$ and $FS_1$, node N2 comprises $RS_3$ and $FS_2$, and node N3 comprises $RS_4$ and $FS_3$; the populations span all three reconstructed types (adaptive RS, non-adaptive RS, and FS), and excitatory (green) and inhibitory (red) projections link populations within and between nodes (legend).
In the top $RS_2$ receives a baseline input with Ornstein–Uhlenbeck fluctuations and an added sinusoidal modulation, while in the bottom $FS_1$ receives a baseline input with Ornstein–Uhlenbeck fluctuations. The three panels show the population firing rates over time for nodes N1, N2 and N3 (colors as in the schematic). Full activation protocol in available on GitHub and reproducible via the corresponding Jupyter Notebook.
